# Monitoring the Antimicrobial Resistance Dynamics of *Salmonella enterica* in Healthy Dairy Cattle Populations at the Individual Farm Level Using Whole-Genome Sequencing

**DOI:** 10.1101/2021.08.20.457169

**Authors:** Laura M. Carroll, Ariel J. Buehler, Ahmed Gaballa, Julie D. Siler, Kevin J. Cummings, Rachel A. Cheng, Martin Wiedmann

**Affiliations:** Department of Food Science, Cornell University, Ithaca, New York, USA; Department of Population Medicine and Diagnostic Sciences, Cornell University, Ithaca, New York, USA

**Keywords:** *Salmonella*, antimicrobial resistance, serotyping, dairy cattle, whole-genome sequencing, evolution, livestock

## Abstract

Livestock represent a possible reservoir for facilitating the transmission of the zoonotic foodborne pathogen *Salmonella enterica* to humans; there is also concern that strains can acquire resistance to antimicrobials in the farm environment. Here, we use whole-genome sequencing (WGS) to characterize *Salmonella* strains (*n* = 128) isolated from healthy dairy cattle and their associated environments on 13 New York State farms to assess the diversity and microevolution of this important pathogen at the level of the individual herd. Additionally, the accuracy and concordance of multiple *in silico* tools are assessed, including: (i) two *in silico* serotyping tools, (ii) combinations of five antimicrobial resistance (AMR) determinant detection tools and one to five AMR determinant databases, and (iii) one antimicrobial minimum inhibitory concentration (MIC) prediction tool. For the isolates sequenced here, *in silico* serotyping methods outperformed traditional serotyping and resolved all un-typable and/or ambiguous serotype assignments. Serotypes assigned *in silico* showed greater congruency with the *Salmonella* whole-genome phylogeny than traditional serotype assignments, and *in silico* methods showed high concordance (99% agreement). *In silico* AMR determinant detection methods additionally showed a high degree of concordance, regardless of the pipeline or database used (≥98% agreement between susceptible/resistant assignments for all pipeline/database combinations). For AMR detection methods that relied exclusively on nucleotide BLAST, accuracy could be maximized by using a range of minimum nucleotide identity and coverage thresholds, with thresholds of 75% nucleotide identity and 50-60% coverage adequate for most pipeline/database combinations. *In silico* characterization of the microevolution and AMR dynamics of each of six serotype groups (*S.* Anatum, Cerro, Kentucky, Meleagridis, Newport, Typhimurium/Typhimurium variant Copenhagen) revealed that some lineages were strongly associated with individual farms, while others were distributed across multiple farms. Numerous AMR determinant acquisition and loss events were identified, including the recent acquisition of cephalosporin resistance-conferring *bla*_CMY_- and *bla*_CTX-M_-type beta-lactamases. The results presented here provide high-resolution insight into the temporal dynamics of AMR *Salmonella* at the scale of the individual farm and highlight both the strengths and limitations of WGS in tracking zoonotic pathogens and their associated AMR determinants at the livestock-human interface.

## 1 Introduction

The foodborne pathogen *Salmonella enterica* is estimated to be responsible for 1.2 million illnesses and 450 deaths each year in the U.S. alone (Scallan et al., 2011). Despite the fact that over 2,600 *Salmonella* serotypes have been described (Issenhuth-Jeanjean et al., 2014), fewer than 100 of these serotypes are responsible for the majority of human infections (Centers for Disease Control and Prevention, 2020). In line with this, some *Salmonella* serotypes may share strong associations with a specific host, an extreme example of which can be seen in the human-restricted nature of *Salmonella* Typhi (Uzzau et al., 2000; Boore et al., 2015). Other serotypes, while not confined exclusively to infection of a single host, may be adapted to a given reservoir; for example, *Salmonella* Choleraesuis, while largely adapted to swine, occasionally infects humans (Uzzau et al., 2000; Chiu et al., 2004).

Cattle are a potential reservoir from which humans can acquire salmonellosis, and infected animals can shed *Salmonella* at irregular intervals for varying periods of time, regardless of whether they express clinical signs of bovine salmonellosis or not (Cummings et al., 2010b; Davidson et al., 2018; Holschbach and Peek, 2018). The bovine reservoir boasts its own repertoire of serotypes that can infect humans, with bovine-associated *Salmonella* serotype Dublin, known for its rare but frequently invasive infections in humans, being arguably the most noteworthy (Taylor et al., 1982; Uzzau et al., 2000; Rodriguez-Rivera et al., 2014; Harvey et al., 2017; Mohammed et al., 2017). However, a range of *Salmonella* serotypes can persist and thrive in cattle, potentially infecting humans via either direct contact with infected animals or through food (Gutema et al., 2019). In a previous survey of 46 dairy cattle herds in New York State, *Salmonella* strains isolated from subclinically infected dairy cattle and associated farm environments spanned 26 serotypes, the most common being Cerro, Kentucky, Typhimurium, Newport, and Anatum (Rodriguez-Rivera et al., 2014). Additionally, antimicrobial-resistant (AMR) isolates were observed on several farms, on numerous occasions, suggesting subclinically infected dairy cattle as a potential source of AMR *Salmonella* (Rodriguez-Rivera et al., 2014).

Numerous studies have employed whole-genome sequencing (WGS) to characterize *Salmonella* from bovine sources (Mather et al., 2013; Agren et al., 2016; Carroll et al., 2017b; Delgado-Suarez et al., 2018; Liao et al., 2019); however, little is known regarding the evolution and AMR acquisition and loss dynamics of *Salmonella* at the single herd/farm level. Furthermore, the bulk of bovine-associated *Salmonella* WGS efforts have focused on clinical veterinary samples and/or epidemic lineages (e.g., *S.* Typhimurium DT104). In this study, 128 non-typhoidal *Salmonella enterica* strains isolated from repeated sampling on 13 New York State dairy cattle farms between 2007 and 2009 were characterized using WGS. All strains were isolated from apparently healthy, subclinically infected bovine hosts, as well as the associated farm environment (Rodriguez-Rivera et al., 2014). Using WGS, we characterized the microevolution of these persistent lineages within each herd, as well as the temporal acquisition and loss of AMR determinants among them. In addition to offering insight into the genomics of *Salmonella* isolated from healthy bovine populations at the individual herd/farm level, we evaluate the accuracy and concordance of multiple *in silico* serotyping and AMR prediction tools. Finally, we provide an in-depth, critical analysis of the strengths and limitations of the methods used here, and we offer guidance to researchers who wish to employ WGS for herd-level pathogen monitoring.

## 2 Materials and Methods

### 2.1 Isolate Selection

*Salmonella enterica* isolates (*n* = 128) obtained from one of 13 dairy farms in New York State were selected to undergo WGS for this study (Supplementary Table S1). All strains were isolated from farms that had undergone surveillance for *Salmonella* for a period of at least 12 months as described previously (Cummings et al., 2010a; Rodriguez-Rivera et al., 2014). Strains were isolated from repeated sampling on each farm between October 2007 and August 2009, from either (i) fecal samples from healthy, subclinically infected dairy cows (referred to hereafter as “bovine” isolates), or (ii) farm environmental swabs (referred to hereafter as “farm environmental” isolates) (Cummings et al., 2010a). All isolates underwent serotyping, phenotypic antimicrobial susceptibility testing, and pulsed-field gel electrophoresis (PFGE) as described previously (Rodriguez-Rivera et al., 2014).

### 2.2 Whole-Genome Sequencing and Data Pre-Processing

Genomic DNA extraction and sequencing library preparation were performed as described previously (Carroll et al., 2017b), and the genomes of all 128 *Salmonella* isolates were sequenced using an Illumina HiSeq platform and 2 x 250 bp paired-end reads. Illumina sequencing adapters and low-quality bases were trimmed using Trimmomatic version 0.33 (using default parameters for Nextera paired-end reads) (Bolger et al., 2014), and FastQC version 0.11.9 (Andrews, 2019) was used to confirm adapter removal and assess read quality. SPAdes version 3.8.0 (Bankevich et al., 2012) was used to assemble genomes *de novo* (using the “careful” option and *k*-mer sizes of 21, 33, 55, 77, 99, and 127), and QUAST version 4.5 (Gurevich et al., 2013) and the “lineage_wf” workflow implemented in CheckM version 1.1.3 (Parks et al., 2015) were used to assess the quality of the resulting assemblies. MultiQC version 1.8 (Ewels et al., 2016) was used to aggregate genome quality metrics. Genome quality statistics are available for all isolates (Supplementary Table S1).

### 2.3 *In Silico* Serotyping

In addition to undergoing traditional serotyping in a laboratory setting (i.e., serological detection of expressed O and H antigens using the White-Kauffmann-Le Minor scheme) as described previously (Rodriguez-Rivera et al., 2014), all 128 assembled *Salmonella* genomes (see section “Whole-Genome Sequencing and Data Pre-Processing” above) underwent *in silico* serotyping using the command line implementations of (i) the *Salmonella In Silico* Typing Resource (SISTR) version 1.0.2 (Yoshida et al., 2016) and (ii) SeqSero2 version 1.1.1 (Zhang et al., 2019) (using SeqSero2’s *k*-mer based workflow). Serotypes assigned using all three methods are available for all 128 isolates (Supplementary Table S1). In cases where a discrepancy existed between the traditional serotype designation and one or more of the *in silico* methods, the serotype assigned using two out of the three methods was selected as the final serotype to be reported (e.g., when assigning strain names to isolates in the manuscript, for phylogeny annotation). To confirm that all serotype assignments were reasonable, a phylogeny was constructed using core single nucleotide polymorphisms (SNPs) detected in all *Salmonella* genomes in this study (see section “Reference-Free SNP Identification and Phylogeny Construction” below).

### 2.4 *In Silico* Antimicrobial Resistance Determinant Detection

Antimicrobial resistance (AMR) determinants were detected in each of the 128 *Salmonella* genomes using five separate approaches: (i) ABRicate (https://github.com/tseemann/abricate) version 0.8 (Seemann, 2018), (ii) AMRFinderPlus version 3.2.3 (Feldgarden et al., 2019), (iii) ARIBA version 2.14.1 (Hunt et al., 2017), (iv) BTyper version 2.3.3 (Carroll et al., 2017a), and (v) SRST2 version 0.2.0 (Inouye et al., 2014). Assembled genomes were used as input for the ABRicate and BTyper approaches, while trimmed Illumina reads were used as input for the SRST2 and ARIBA approaches. Prokka version 1.12 (Seemann, 2014) was used to annotate each assembled genome, and the resulting GFF (.gff) and FASTA (.faa and .ffn) files were used as input for the AMRFinderPlus approach. For the ABRicate approach, the following AMR gene databases were tested (each accessed June 11, 2018 via ABRicate’s abricate-get_db command): (i) the Antibiotic Resistance Gene-ANNOTation database (ARG-ANNOT) (Gupta et al., 2014), (ii) the Comprehensive Antibiotic Resistance Database (CARD) (Jia et al., 2017), (iii) the National Center for Biotechnology Information’s (NCBI’s) Bacterial Antimicrobial Resistance Reference Gene Database (NCBI) (Feldgarden et al., 2019), and (iv) the ResFinder database (ResFinder) (Zankari et al., 2012). For each genome and database combination, minimum AMR gene identity and coverage thresholds ranging from 50-100% (5% increments) and 0-100% (10% increments) were tested, respectively. For the BTyper approach, the (i) ARG-ANNOT v3 and (ii) MEGARes version 1.0.1 (Lakin et al., 2017) databases available with BTyper version 2.3.3 were used, with the minimum AMR gene identity and coverage thresholds varied in a manner identical to the ABRicate approach. For the SRST2 approach, the (i) ARG-ANNOT and (ii) ResFinder databases available with SRST2 version 0.2.0 were tested, using default thresholds. For the ARIBA approach, the following databases were tested (each accessed June 13, 2019 using ARIBA’s getref command): (i) the version of ARG-ANNOT available with SRST2, (ii) CARD, (iii) MEGARes, (iv) NCBI, and (v) ResFinder, with all default thresholds used. For the AMRFinder approach, the latest version of the AMRFinder database was used (accessed December 6, 2019), along with the organism-specific database for *Salmonella*.

### 2.5 *In Silico* Prediction of Antimicrobial Minimum Inhibitory Concentration Values

The PATRIC3 antimicrobial minimum inhibitory concentration (MIC) prediction model for *Salmonella* (Nguyen et al., 2019) (accessed June 13, 2019) was used to predict MIC values for each of the 128 *Salmonella* isolates in this study, using the assembled genome of each as input. The following were used as dependencies: kmc version 3.0 (Kokot et al., 2017) and XGBoost version 0.82 (Chen and Guestrin, 2016).

### 2.6 Prediction of Phenotypic Susceptible-Intermediate-Resistant Classifications Using *In Silico* Methods

All 128 *Salmonella* isolates underwent phenotypic antimicrobial susceptibility testing with a panel of 15 antimicrobials (i.e., amikacin, amoxicillin-clavulanic acid, ampicillin, cefoxitin, ceftiofur, ceftriaxone, chloramphenicol, ciprofloxacin, gentamicin, kanamycin, nalidixic acid, streptomycin, sulfamethoxazole-trimethoprim, sulfisoxazole, and tetracycline) using the Sensititre® system (Trek Diagnostic Systems Ltd., Cleveland, OH) available at Cornell University’s Animal Health Diagnostic Center as described previously (Rodriguez-Rivera et al., 2014). A “true” (i.e., phenotypic) susceptible-intermediate-resistant (SIR) classification for each of the 15 antimicrobials was obtained for 126 *Salmonella* isolates by comparing raw MIC values to NARMS breakpoints for *Salmonella* (accessed March 23, 2020; Supplementary Table S1). For streptomycin, the 1996-2013 NARMS breakpoints were used, as this was compatible with the concentrations used at the time of phenotypic testing (Rodriguez-Rivera et al., 2014). For sulfisoxazole, isolates with MIC > 256 were classified as resistant, as a concentration of 512 μg/mL was not tested. While raw MIC values were unavailable for two isolates (BOV_KENT_16_04-03-08_R8-0967 and ENV_MELA_01_01-10-08_R8-0165; Supplementary Table S1), both isolates had previously been categorized as pan-susceptible to all 15 antimicrobials (a classification that was maintained here, as all *in silico* methods correctly classified these isolates as pan-susceptible).

Supplemental Table S4 of the AMRFinder validation paper (Feldgarden et al., 2019) was used to identify known AMR determinant/phenotype associations for AMR determinants detected by each of the AMR determinant detection pipeline/database combinations described above (see section “*In Silico* Antimicrobial Resistance Determinant Detection”) and all of the 15 antimicrobials tested in this study (Supplementary Table S2). If a detected AMR gene was not identified in the AMRFinder literature search, the gene name was queried in CARD, and the resulting literature linked to the CARD entry was searched for known AMR genotype/phenotype associations (Supplementary Table S2). An isolate was predicted to be resistant to a particular antimicrobial if it possessed one or more AMR determinants known to confer resistance to that antimicrobial; if it did not possess any AMR determinants known to confer resistance to that antimicrobial, the isolate was predicted to be susceptible to that antimicrobial (Supplementary Table S2). For each AMR determinant detection pipeline/database combination, the caret package (Kuhn, 2008) in R version 3.6.1 (R Core Team, 2019) was used to construct a confusion matrix and calculate accuracy scores, Cohen’s kappa coefficients, and other statistics (Supplementary Table S3) by treating “true” susceptible/resistant classifications obtained using phenotypic susceptibility testing as a reference. Cases of intermediate phenotypic resistance were treated as susceptible, as it resulted in slightly higher accuracy scores for all pipeline/database combinations for this particular data set. Because *in silico* prediction of susceptibility/resistance was highly dependent on prior knowledge of AMR determinants and the antimicrobials to which they conferred resistance, the concordance of all pipeline/database combinations was assessed by comparing each pipeline/database combination to results obtained using the SRST2 pipeline/ARG-ANNOT database combination.

To assess the ability of the MIC prediction method implemented in PATRIC3 to predict *Salmonella* SIR classification (see section “*In Silico* Prediction of Antimicrobial Minimum Inhibitory Concentration Values” above), predicted MIC values for 14 antimicrobials produced using PATRIC3 were used to predict the SIR status of each of the 128 *Salmonella* isolates using the same NARMS breakpoints used for phenotypic testing. Azithromycin MICs produced by PATRIC3 were excluded, as azithromycin was not among the 15 antimicrobials used here for phenotypic testing. The ability of PATRIC3 to predict amikacin resistance was also not evaluated, as amikacin is not among the antimicrobials queried by PATRIC3. A confusion matrix was constructed as described above, using predicted SIR classifications derived from predicted MIC values produced by PATRIC3 and NARMS breakpoints. Additionally, the deviation of raw MIC predictions produced by PATRIC3 (*MIC_PATRIC3_*) from “true” raw MIC predictions produced using phenotypic testing (*MIC_Phenotypic_*) in number of dilution factors (*N_dilution factors_*) was assessed using the following equation:

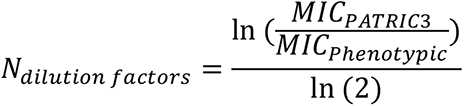

where ln corresponds to the natural logarithm. For example: if PATRIC3 predicted an MIC value of 8 and the “true” MIC value obtained with phenotypic testing was 2, then ln(8/2)/ln(2) = 2; this means that the PATRIC3 prediction of 8 is +2 dilution factors away from the “true” MIC of 2 (as dilution used for MIC are 2 fold serial dilutions, e.g., 2 μg/mL, 4 μg/mL, 8 μg/mL).

### 2.7 Re-testing of Isolates with Highly Incongruent AMR Phenotypes

Several (*n* = 21) isolates possessed a phenotypic AMR SIR profile which was deemed to be highly incongruent with its predicted *in silico* AMR profile, regardless of the *in silico* pipeline/database used (Supplementary Table S4). For example, *S.* Cerro isolate BOV_CERO_35_10−02−08_R8−2685 was resistant to nine antimicrobials but did not harbor any known acquired AMR genes (Supplementary Table S4). Similarly, *S.* Newport isolate ENV_NEWP_62_03−05−09_R8−3442 itself was pan-susceptible, but harbored multiple acquired AMR genes (e.g., *bla*_CMY-2_, *floR*, *sul2*, *tetA*), which conferred multidrug resistance in closely related *S.* Newport isolates (Supplementary Table S4). To address these incongruencies, 21 selected *Salmonella* isolates underwent phenotypic antimicrobial susceptibility re-testing (conducted September 16, 2020) as described above (see section “Prediction of Phenotypic Susceptible-Intermediate-Resistant Classifications Using *In Silico* Methods”), with the exception of amikacin and kanamycin, as the contemporary panel did not include these antimicrobials (Supplementary Table S4).

Kanamycin testing was conducted separately using a gradient diffusion assay (Jorgensen and Ferraro, 2009) according to the manufacturer’s instructions (BioMérieux Kanamycin Strip KM 256, product number 412381). Briefly, bacterial isolates were streaked for single colonies onto Brain Heart Infusion (BHI, Becton Dickinson [BD], Franklin Lakes, NJ, USA) agar plates from frozen glycerol stocks. Precultures were prepared by inoculating a single colony in 3 mL Mueller-Hinton (MH) broth (BD Difco), followed by incubating at 37°C with shaking at 200 rpm for 12-14h. The precultures were used to inoculate tubes with 5 mL MH broth at 1:200 dilution, and the tubes were incubated at 37°C with shaking at 200 rpm for 5 hours. Four mL of melted MH soft agar medium (0.7% agar) were mixed with 100 μL of culture and poured onto Petri plates containing 15 mL of MH agar medium (0.7% agar), and the plates were dried for 5 min. Kanamycin gradient strips were laid on top of the soft agar, and the plates were incubated at 35°C for 18 hours. MIC values were determined by evaluating the inhibition zone using a magnifying lens according to the manufacturer’s instructions.

MIC values obtained from re-testing these isolates were interpreted within NARMS breakpoints as described above (see section “Prediction of Phenotypic Susceptible-Intermediate-Resistant Classifications Using *In Silico* Methods”) and are reported in the main manuscript (with the exception of amikacin; due to its exclusion from the contemporary panel, original MIC values are reported). Original and updated MIC and SIR values for all 21 isolates are available in Supplementary Table S4.

### 2.8 *In Silico* Plasmid Replicon Detection

Plasmid replicons were detected in all *Salmonella* genome assemblies using ABRicate and the PlasmidFinder database (accessed June 11, 2018 via ABRicate’s abricate-get_db command). For a plasmid replicon to be considered present in a genome, minimum nucleotide BLAST (blastn) (Camacho et al., 2009) identity and coverage values of 80 and 60%, respectively, were used (Carattoli et al., 2014).

### 2.9 Reference-Free SNP Identification and Phylogeny Construction

A reference-free approach was used to compare the 128 *Salmonella* genomes sequenced in this study to 442 of the 445 *Salmonella* genomes described by Worley et al. (Worley et al., 2018); three genomes were omitted because their Sequence Read Archive (SRA) data was not publicly available at the time of access (February 20, 2019). Raw reads for each of the 442 publicly available genomes were downloaded from SRA (Leinonen et al., 2011; Kodama et al., 2012) and processed and assembled as described above (see section “Whole-Genome Sequencing and Data Pre-Processing” described above). kSNP3 version 3.1 (Gardner and Hall, 2013; Gardner et al., 2015) was used to identify core SNPs among all 570 assembled *Salmonella* genomes, using the optimal *k-*mer size determined by Kchooser (*k* = 19). IQ-TREE version 1.6.10 (Nguyen et al., 2015) was used to construct a maximum likelihood (ML) phylogeny using the resulting core SNPs and the optimal nucleotide substitution model identified using ModelFinder (determined using model Bayesian Information Criteria [BIC] values); the optimal model was the transversion model with unequal, empirical base frequencies, an ascertainment bias correction, and the FreeRate model with six categories, i.e., the TVM +F+ASC+R6 model (Yang, 1995; Lewis, 2001; Soubrier et al., 2012; Kalyaanamoorthy et al., 2017). Bootstrapping was performed using 1,000 replicates of the Ultrafast Bootstrap method (Minh et al., 2013; Hoang et al., 2018). The resulting ML phylogeny was annotated in R using the bactaxR package (Carroll et al., 2020b) and the following dependencies: ape (Paradis et al., 2004; Paradis and Schliep, 2019), dplyr (Wickham et al., 2020), ggtree (Yu et al., 2017; Yu et al., 2018), phylobase (R Hackathon, 2019), phytools (Revell, 2012), and reshape2 (Wickham, 2007).

### 2.10 Pan-Genome Characterization

GFF files produced by Prokka (see section “*In Silico* Antimicrobial Resistance Determinant Detection” above) were used as input for Roary version 3.12.0 (Page et al., 2015), which was used to identify orthologous gene clusters at a 70% protein BLAST (blastp) identity threshold. The resulting gene presence/absence matrix produced by Roary was used as input for besPLOT (https://github.com/lmc297/besPLOT) (Carroll et al., 2020a), which was used to perform non-metric multidimensional scaling (NMDS) (Kruskal, 1964) and construct plots in two dimensions using a Jaccard distance metric and the following dependencies in R: vegan version 2.5-6 (Oksanen et al., 2019), shiny version 1.4.0.2 (Chang et al., 2020), ggplot2 version 3.3.0 (Wickham, 2016), plyr version 1.8.6 (Wickham, 2011), dplyr version 0.8.5 (Wickham et al., 2020), cluster version 2.1.0 (Maechler et al., 2019), and ggrepel version 0.8.2 (Slowikowski, 2020).

Clustering based on gene presence/absence was assessed for each of the following grouping factors: (i) serotype, (ii) farm, and (iii) isolation source (i.e., bovine or farm environmental). For each of the three grouping factors, the following three statistical tests were performed, using the gene presence/absence matrix produced by Roary, a Jaccard distance metric, and 10,000 permutations: (i) the permutest and betadisper functions in R’s vegan package (Oksanen et al., 2019) were used to conduct an ANOVA-like permutation test (Anderson, 2006) to test if group dispersions were homogenous (referred to hereafter as the PERMDISP2 test); (ii) analysis of similarity (ANOSIM) (Clarke, 1993) using the anosim function in the vegan package in R was used to determine if the average of the ranks of within-group distances was greater than or equal to the average of the ranks of between-group distances (Anderson and Walsh, 2013); (iii) permutational analysis of variance (PERMANOVA) (Anderson, 2001) using the adonis2 function in the vegan package in R was used to determine if group centroids were equivalent. For all tests, a Bonferroni correction was applied to correct for multiple comparisons.

Potential clustering based on AMR gene presence/absence was additionally assessed for the same three grouping factors (serotype, farm, and isolation source), using the presence and absence of AMR determinants detected by AMRFinderPlus as input (i.e., AMR and stress response determinants identified using the “plus” option in AMRFinderPlus). All steps were performed as described above, and a Bonferroni correction was used to correct for multiple comparisons.

### 2.11 Reference-Based Core SNP Identification Within Serotypes

For each individual serotype, core SNPs were identified among genomes assigned to that serotype using a reference-based approach. For each serotype, Snippy version 4.3.6 (https://github.com/tseemann/snippy) (Seemann, 2019b) was used to identify core SNPs among all representatives assigned to the serotype, using the trimmed Illumina paired-end reads of each genome as input (see section “Whole-Genome Sequencing and Data Pre-Processing” above), one of six high-quality assembled genomes from isolates in this study as a reference genome (Supplementary Table S1), and the following dependencies: BWA MEM version 0.7.13-r1126 (Li and Durbin, 2009; Li, 2013), Minimap2 version 2.15 (Li, 2018), SAMtools version 1.8 (Li et al., 2009), BEDtools version 2.27.1 (Quinlan and Hall, 2010; Quinlan, 2014), BCFtools version 1.8 (Li, 2011), FreeBayes version v1.1.0-60-gc15b070 (Garrison and Marth, 2012), vcflib version v1.0.0-rc2 (Cleary et al., 2015), vt version 0.57721 (Tan et al., 2015), SnpEff version 4.3T (Cingolani et al., 2012), samclip version 0.2 (Seemann, 2019a), seqtk version 1.2-r102-dirty (Li, 2019), and snp-sites version 2.4.0 (Page et al., 2016). Gubbins version 2.3.4 (Croucher et al., 2015) was used to identify and remove recombination within the full alignment that resulted, and the filtered alignment produced by Gubbins was queried using snp-sites to produce an alignment of core SNPs for each serotype.

### 2.12 Construction of Within-Serotype Phylogenies

For each serotype, IQ-TREE version 1.6.10 was used to construct a ML phylogeny, using core SNPs detected among all isolates assigned to the serotype as input (see “Reference-Based Core SNP Identification Within Serotypes” section above), the optimal ascertainment bias-aware nucleotide substitution model selected using ModelFinder, and 1,000 replicates of the UltraFast bootstrap approximation. The temporal structure of each resulting ML phylogeny was assessed using the *R*^2^ value produced by the best-fitting root in TempEst version 1.5.1 (Supplementary Table S5) (Rambaut et al., 2016).

A tip-dated phylogeny was then constructed for each serotype using BEAST version 2.5.0 (Bouckaert et al., 2014; Bouckaert et al., 2019), the appropriate core SNP alignment, an initial clock rate of 2.79×10^−7^ substitutions/site/year (Leekitcharoenphon et al., 2016), and an ascertainment bias correction to account for the use of solely variant sites (Bouckaert, 2014). The modelTest function in R’s phangorn package (Schliep, 2011) was used to select an appropriate nucleotide substitution model for each serotype, and the implementation of the model with the lowest BIC value in BEAST 2’s SSN package (Bouckaert and Xie, 2017) was selected. For each serotype, combinations of either a strict or lognormal relaxed molecular clock (Drummond et al., 2006) and either a coalescent constant or coalescent Bayesian Skyline (Drummond et al., 2005) population model were tested. For all models, a lognormal prior was used for the clockRate/ucldMean parameters (in real space, M = 2.058×10^−6^ and S = 2.0, yielding a median value of 2.79×10^−7^ substitutions/site/year, and 2.5% and 97.5% quantiles of 5.53×10^−9^ and 1.40×10^−5^ substitutions/site/year, respectively). For each molecular clock and population model combination for each serotype, three independent stepping stone sampling analyses (Xie et al., 2011) were performed with BEAST 2, using 10 steps of at least 100 million generations. Bayes factors (Kass and Raftery, 1995) were calculated to determine the optimal combination of molecular clock and population model for each serotype (Supplementary Table S5). Ten independent BEAST 2 runs were then performed using the optimal model for each serotype, using chain lengths of at least 100 million generations, sampling every 10,000 generations. Tracer version 1.7.1 (Rambaut et al., 2018) was used to ensure adequate mixing of each independent run. The resulting log and tree files were aggregated using LogCombiner-2, and TreeAnnotator-2 (Heled and Bouckaert, 2013) was used to produce a maximum clade credibility tree using 10% burn-in. The phylogenies were annotated using R and the following packages: ggplot2, ggtree, and phylobase. Bayesian skyline plots for all serotype groups analyzed using BEAST2 are also available (Supplementary Figure S1); however, due to the limited number of available isolates surveyed among each serotype and the short temporal range queried here, potential changes in effective population sizes may not be robust and are thus not discussed in this study.

### 2.13 Data Availability

Illumina reads are available for all isolates sequenced in this study under NCBI Bioproject Accession PRJNA756552. NCBI BioSample accession numbers for each individual isolate, as well as all associated metadata and genome quality statistics, are available in Supplementary Table S1. All BEAST 2 XML files used for temporal phylogeny construction are available at https://github.com/lmc297/zru_farms.

## 3 Results

### 3.1 *In silico* serotyping of bovine-associated *Salmonella* resolves incongruencies between traditional serotyping and whole-genome phylogeny

A total of 128 *Salmonella* strains isolated from healthy (i.e., subclinically infected) dairy cattle (*n* = 39) and their associated farm environments (*n* = 89) on 13 different New York State farms underwent WGS (Supplementary Table S1). In addition to undergoing traditional serotyping in a laboratory setting, all isolates were assigned serotypes *in silico* using both (i) SISTR and (ii) SeqSero2 (Supplementary Table S1). Importantly, serotypes assigned *in silico* using SISTR and/or SeqSero2 were able to resolve all un-typable and/or ambiguous serotypes assigned using traditional serotyping (Supplementary Table S1). Furthermore, *in silico* serotypes assigned using (i) SISTR’s core-genome multi-locus sequence typing (cgMLST) approach and (ii) SeqSero2 were both highly congruent with the *Salmonella* whole-genome phylogeny (Figure 1) and highly concordant with each other: 127 of 128 (99.2%) *Salmonella* isolates sequenced in this study were assigned to identical *in silico* serotypes using both SISTR cgMLST and SeqSero2 (Supplementary Table S1), with 100% concordance observed for six of seven observed *in silico* serotype groups (i.e., *S.* Anatum, *S.* Cerro, *S.* Meleagridis, *S.* Minnesota, *S.* Newport, and *S.* Typhimurium and its variants, assigned to *n* = 15, 13, 20, 1, 16, and 27 isolates, respectively). Among *S.* Kentucky (*n* = 36), a single incongruent isolate was observed (ENV_KENT_16_12-04-07_R8-0061), as SeqSero2 could not detect an O-antigen within the genome and was thus unable to assign this isolate to any serotype. This isolate was assigned a serotype of 8,20:-:z6 using traditional serotyping (*S.* Kentucky has antigenic formula 8,20:i:z6); SISTR classified the isolate as *S.* Kentucky, and the isolate clustered among the *S.* Kentucky isolates sequenced in this study within the *Salmonella* whole-genome phylogeny (Figure 1 and Supplementary Table S1).

**Figure 1.**
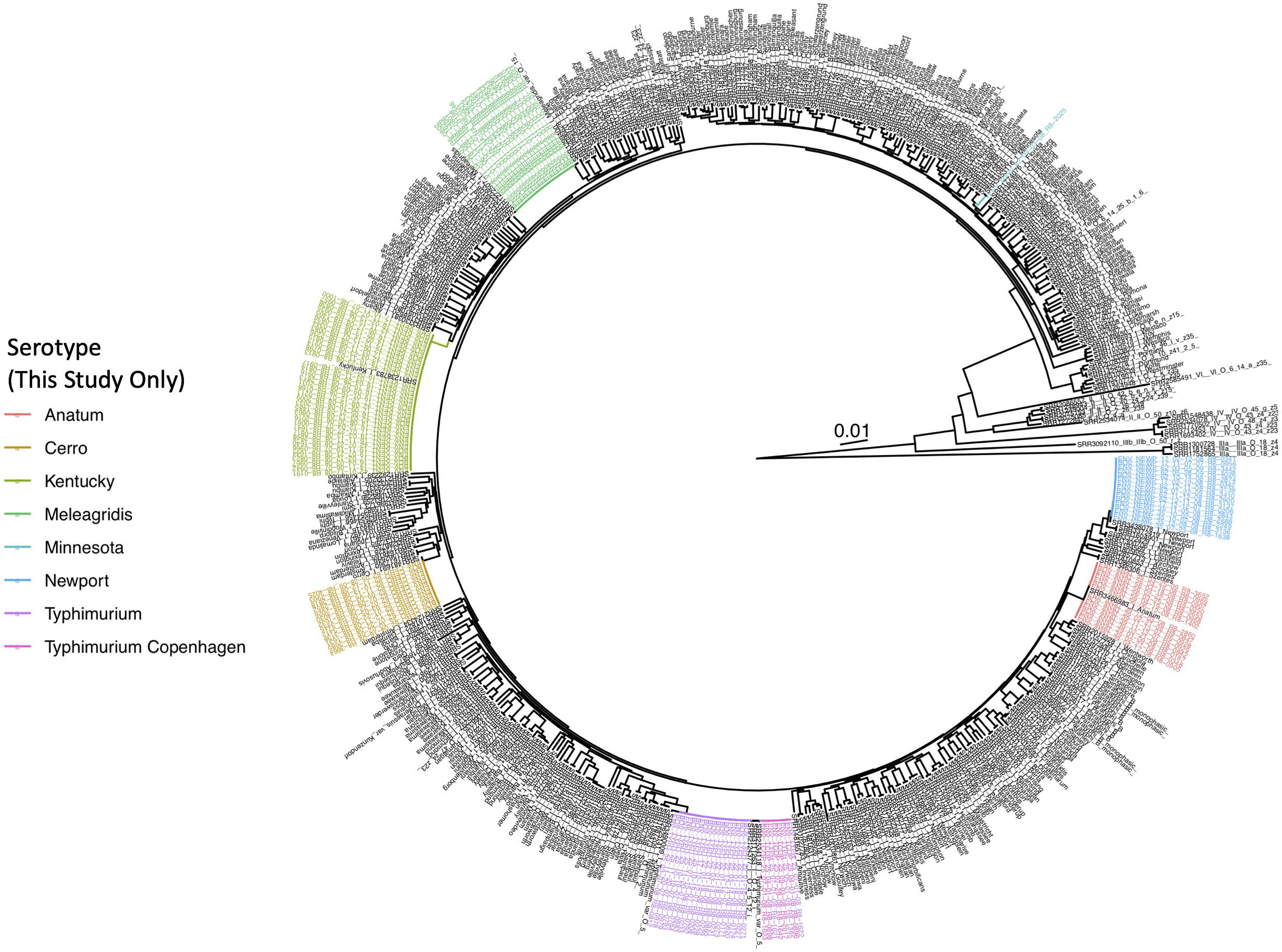
Maximum likelihood phylogeny constructed using core SNPs identified among 570 *Salmonella* isolate genomes. Publicly available genomes are denoted by black tip labels (*n* = 442), while genomes of bovine- and bovine farm-associated strains isolated in conjunction with this study are denoted by colored tip labels (*n* = 128). In cases where a discrepancy existed between the traditional serotype designation of an isolate and one or more *in silico* methods (i.e., SISTR and SeqSero2), the serotype assigned using two out of the three methods was selected as the final serotype to be used for phylogeny annotation. The phylogeny is rooted at the midpoint with branch lengths reported in substitutions per site. Core SNPs were identified among all genomes using kSNP3, while the phylogeny was constructed and annotated using IQ-TREE and bactaxR/ggtree, respectively.

When variants of the *S.* Typhimurium serotype (*n* = 27) were considered, discrepancies were observed between traditional serotype assignments and both *in silico* methods (Supplementary Table S1). While SeqSero2 could differentiate between *S.* Typhimurium and the O5-variant of *S.* Typhimurium (also known as *S.* Typhimurium variant Copenhagen; “*S.* Typhimurium Copenhagen” is used hereafter), SISTR was unable to differentiate the two (Supplementary Table S1), as noted previously (Ibrahim and Morin, 2018; Zhang et al., 2019). However, *S.* Typhimurium and *S.* Typhimurium Copenhagen serotype assignments obtained using SeqSero2 and traditional serotyping did not always agree, as five of 27 *S.* Typhimurium/*S.* Typhimurium Copenhagen assignments (18.5%) differed between the two methods (Supplementary Table S1). For four of the five incongruent isolates, SeqSero2 assigned an isolate to *S.* Typhimurium Copenhagen, while traditional serotyping assigned a serotype of *S.* Typhimurium; for one isolate, the opposite scenario applied (Supplementary Table S1). Furthermore, the lineages formed by isolates classified here as *S.* Typhimurium Copenhagen using either traditional serotyping or SeqSero2, as well as two *S.* Typhimurium Copenhagen genomes from a previous study (Worley et al., 2018), were polyphyletic (Figure 1); consequently, the whole-genome phylogeny could not be used to reliably differentiate these two variants.

For the remainder of this study, serotypes assigned consistently with at least two out of the three methods (i.e., traditional serotyping, SeqSero2, and SISTR cgMLST) were selected as the final serotype to be reported for each isolate. Nine of the 13 farms surveyed here harbored *Salmonella* isolates that belonged to a single serotype, while two farms harbored two serotypes or serotype variants (Farms 25 and 35 harbored Typhimurium/Typhimurium Copenhagen and Cerro/Newport, respectively; Supplementary Table S1). The remaining two farms harbored three *Salmonella* serotypes (Farms 17 and 62 harbored Kentucky/Newport/Typhimurium and Cerro/Minnesota/Newport, respectively; Supplementary Table S1).

### 3.2 *In silico* methods predict antimicrobial susceptibility and resistance among bovine-associated *Salmonella* with high accuracy and concordance

Using a 15-antimicrobial panel and NARMS breakpoints for *Salmonella*, more than half of all isolates in this study (81 of 128; 63.3%) were classified as susceptible to all 15 antimicrobials tested, while 38 isolates (29.7%) were classified as resistant to two or more antimicrobials (obtained after the 15-antimicrobial panel was re-run for 22 isolates to resolve discrepancies between *in silico* predictions and phenotypic AMR data; Supplementary Tables S1 and S4).

Regardless of choice of AMR determinant detection pipeline and AMR determinant database, all pipeline/database combinations performed nearly identically when given the task of predicting phenotypic AMR susceptibility/resistance to 15 antimicrobials using known AMR determinant-phenotype associations (Figure 2, Table 1, and Supplementary Tables S2 and S3). Furthermore, all pipeline/database combinations showed an extremely high degree of concordance (98.0% or greater for all pipeline/database combinations; Supplementary Figure S2). The overall accuracy of all *in silico* AMR determinant detection pipeline/database combinations ranged from 95.8-97.4%, with the SRST2 AMR detection tool/ARG-ANNOT AMR determinant database combination achieving the highest accuracy for this data set (Figure 2, Table 1, and Supplementary Table S3). The ARIBA/CARD pipeline/database combination achieved the highest specificity, although all pipeline/database combinations were able to predict phenotypic AMR with high specificity (>99.0%; Figure 2, Table 1, and Supplementary Table S3). Sensitivity ranged from 71.8-84.4%, with SRST2 achieving the highest sensitivities (84.4 and 84.0% for the ARG-ANNOT and ResFinder databases, respectively; Table 1 and Supplementary Table S3).

**Figure 2.**
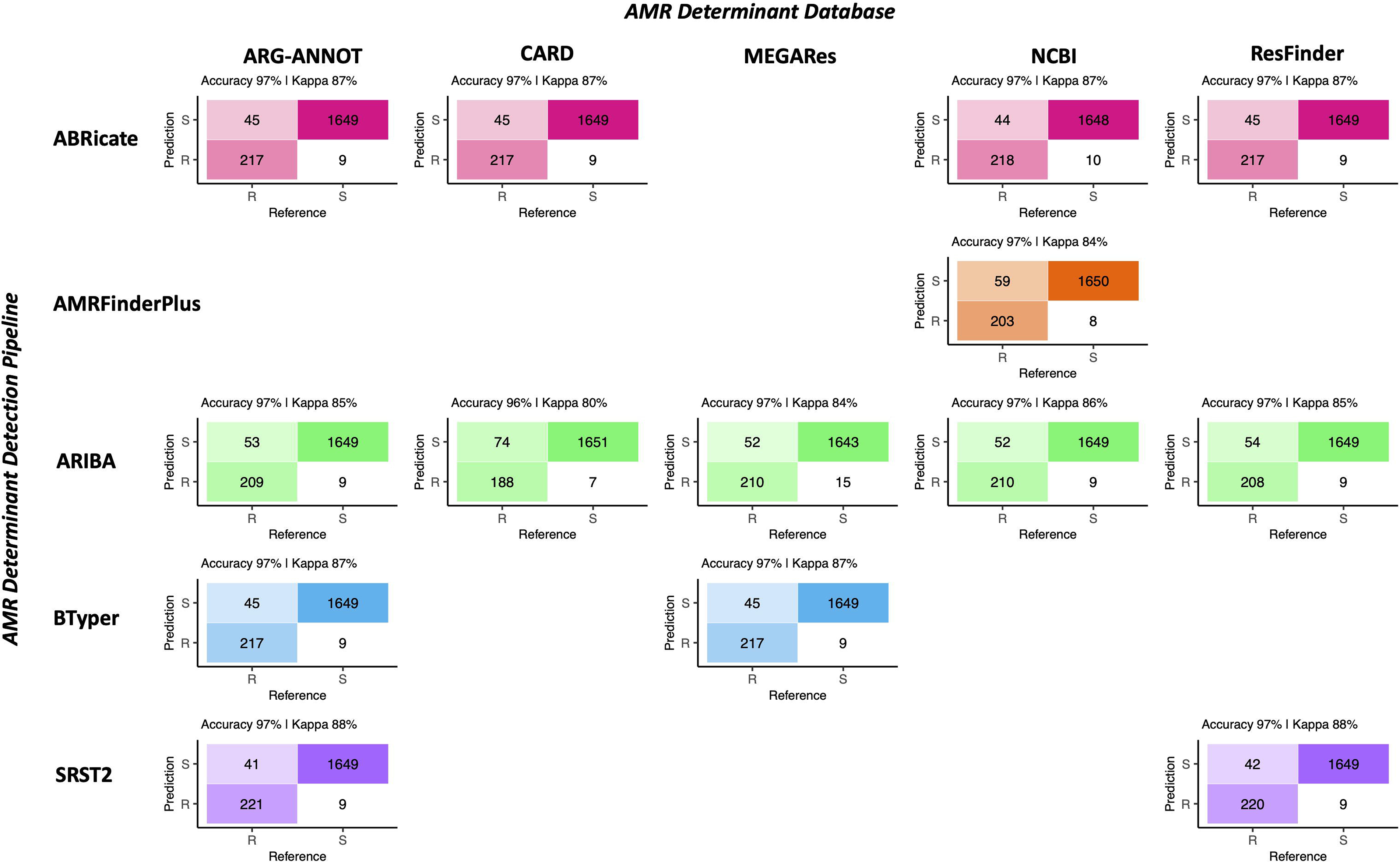
Confusion matrices showcasing agreement between susceptible-intermediate-resistant (SIR) classification of 128 *Salmonella* isolates obtained using phenotypic resistance testing (denoted as the matrix “Reference”) and five *in silico* methods (denoted as the matrix “Prediction”) for 15 antimicrobials. Accuracy values above each matrix denote the percentage of correctly classified instances out of all instances. Kappa values above each matrix denote Cohen’s kappa coefficient for the matrix, reported as a percent. For the phenotypic method, SIR classification was determined using NARMS breakpoints for *Salmonella* (accessed March 23, 2020). For the *in silico* antimicrobial resistance (AMR) determinant detection approaches, combinations of five pipelines and one to five AMR determinant databases were tested; isolate genomes that harbored one or more AMR determinants previously known to confer resistance to a particular antimicrobial were categorized as resistant to that antimicrobial (“R”), while those which did not were categorized as susceptible (“S”; see Supplementary Table S2 for all detected AMR determinants and their associated resistance classifications). For all AMR determinant detection methods, isolates that showed intermediate phenotypic resistance to an antimicrobial were categorized as susceptible (“S”) rather than resistant, as this classification produced slightly better accuracy scores for all pipeline/database combinations. For AMR determinant detection methods that relied on nucleotide BLAST (i.e., ABRicate and BTyper), the confusion matrix obtained using the nucleotide identity and coverage threshold combination that produced the highest accuracy are shown (Supplementary Figure S3); for all other methods, confusion matrices obtained using default detection parameters are shown.

**Table 1.** Statistics for the 15-antimicrobial phenotypic susceptibility/resistance prediction task for all antimicrobial resistance (AMR) determinant pipeline/database combinations.^a^

| AMR Pipeline | AMR Database | % Accuracy (95% Confidence Interval) | Cohen's Kappa (%) | Corrected Accuracy *P*-Value<sup>b</sup> | Corrected McNemar's Test *P*-Value<sup>b</sup> | Sensitivity (%) | Specificity (%) |
| --- | --- | --- | --- | --- | --- | --- | --- |
| ABRicate | ARG-ANNOT | 97.2 (96.3-97.9) | 87.3 | 1.72E-59 | 2.67E-05 | 82.8 | 99.5 |
| ABRicate | CARD | 97.2 (96.3-97.9) | 87.3 | 1.72E-59 | 2.67E-05 | 82.8 | 99.5 |
| ABRicate | NCBI | 97.2 (96.3-97.9) | 87.4 | 1.72E-59 | 9.94E-05 | 83.2 | 99.4 |
| ABRicate | ResFinder | 97.2 (96.3-97.9) | 87.3 | 1.72E-59 | 2.67E-05 | 82.8 | 99.5 |
| AMRFinder Plus | NCBI | 96.5 (95.6-97.3) | 83.9 | 1.41E-50 | 1.41E-08 | 77.5 | 99.5 |
| ARIBA | ARG-ANNOT | 96.8 (95.9-97.5) | 85.3 | 7.37E-54 | 6.63E-07 | 79.8 | 99.5 |
| ARIBA | CARD | 95.8 (94.8-96.6) | 79.9 | 3.08E-42 | 3.14E-12 | 71.8 | 99.6 |
| ARIBA | MEGARes | 96.5 (95.6-97.3) | 84.3 | 1.41E-50 | 1.53E-04 | 80.2 | 99.1 |
| ARIBA | NCBI | 96.8 (95.9-97.6) | 85.5 | 1.55E-54 | 1.06E-06 | 80.2 | 99.5 |
| ARIBA | ResFinder | 96.7 (95.8-97.5) | 85.0 | 3.45E-53 | 4.15E-07 | 79.4 | 99.5 |
| BTyper | ARG-ANNOT | 97.2 (96.3-97.9) | 87.3 | 1.72E-59 | 2.67E-05 | 82.8 | 99.5 |
| BTyper | MEGARes | 97.2 (96.3-97.9) | 87.3 | 1.72E-59 | 2.67E-05 | 82.8 | 99.5 |
| SRST2 | ARG-ANNOT | 97.4 (96.6-98.1) | 88.4 | 1.69E-62 | 1.63E-04 | 84.4 | 99.5 |
| SRST2 | ResFinder | 97.3 (96.5-98.0) | 88.1 | 9.85E-62 | 1.04E-04 | 84.0 | 99.5 |

<sup>a</sup>Statistics were calculated using the confusionMatrix function in the caret package in R, with resistant (“R”) phenotypes/genotypes treated as the “positive” result and susceptible (“S”) phenotypes/genotypes treated as the “negative” result; See Supplementary Table S3 for an extended version of this table.
<sup>b</sup>Adjusted using a Bonferroni correction.

For the AMR determinant detection pipelines that relied on nucleotide BLAST (i.e., ABRicate and BTyper), a range of minimum percent nucleotide identity and coverage thresholds were additionally tested (i.e., all combinations of 50-100% nucleotide identity in increments of 5% and 0-100% coverage in increments of 10%; Supplementary Figure S3) so that the optimal combination(s) could be established for the isolate genomes sequenced here. For ABRicate/ARG-ANNOT, ABRicate/NCBI, and ABRicate/ResFinder, maximum accuracy was achieved using minimum coverage thresholds of 60, 50, and 50-60%, respectively, and 75-95% nucleotide identity thresholds (Supplementary Figure S3). For ABRicate/CARD, minimum thresholds of 60% coverage and 75% nucleotide identity were optimal (Supplementary Figure S3). For BTyper/ARG-ANNOT, maximum accuracy was achieved using 60% coverage and 50-95% nucleotide identity; for BTyper/MEGARes, 50-60% coverage and 95% nucleotide identity were the optimal thresholds (Supplementary Figure S3).

The performance of the PATRIC3 *in silico* MIC prediction method was additionally evaluated (Figure 3 and Supplementary Figure S4). PATRIC3 was able to correctly classify *Salmonella* isolates as susceptible-intermediate-resistant (SIR) based on NARMS breakpoints with an overall accuracy of 92.9% (95% confidence interval 91.6-94.1%, accuracy *P*-value [accuracy > no information rate] < 1.25E-26; Figure 3). At the individual antimicrobial level, PATRIC3 achieved >90% SIR prediction accuracy for 12 of 14 antimicrobials; only sulfisoxazole and tetracycline resistance prediction accuracies were < 90% (83.6 and 68.0%, respectively; Figure 3 and Supplementary Figure S4).

**Figure 3.**
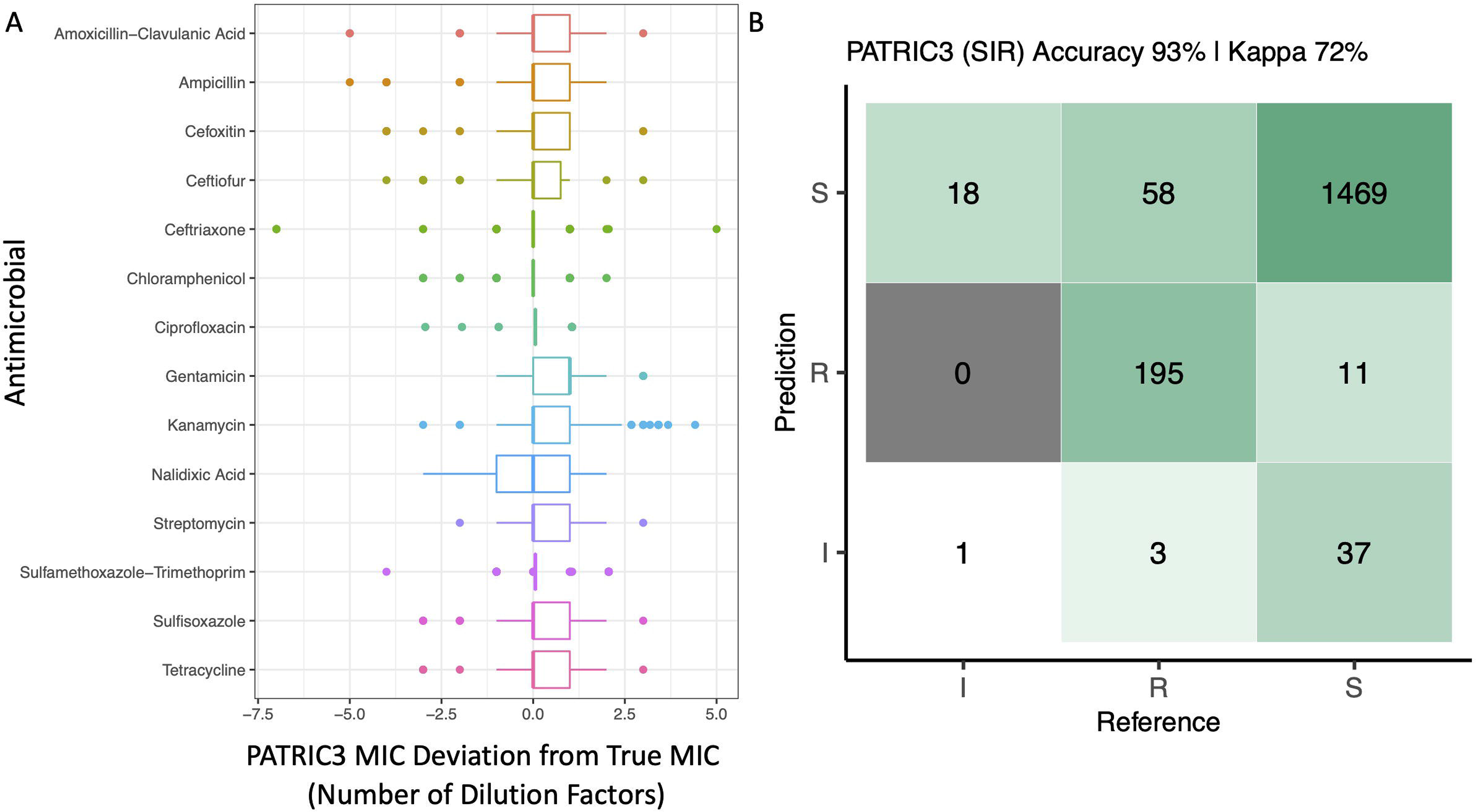
(A) Deviation of PATRIC3 predicted minimum inhibitory concentration (MIC) values from “true” MIC values obtained using phenotypic resistance testing for 14 antimicrobials (Y-axis), reported in number of dilution factors (X-axis). For each box plot, lower and upper box hinges correspond to the first and third quartiles, respectively. Lower and upper whiskers extend from the hinge to the smallest and largest values no more distant than 1.5 times the interquartile range from the hinge, respectively. Points represent pairwise distances that fall beyond the ends of the whiskers. Only isolates with raw MIC values available are included (126 of 128 isolates). (B) Confusion matrix showcasing agreement between susceptible-intermediate-resistant (SIR) classification of all 128 *Salmonella* isolates obtained using phenotypic resistance testing (denoted as the matrix “Reference”) and PATRIC3 (denoted as the matrix “Prediction”) for 14 antimicrobials. The accuracy value above the matrix denotes the percentage of correctly classified instances out of all instances. The Kappa value above the matrix denotes Cohen’s kappa coefficient for the matrix, reported as a percent. For both the phenotypic and PATRIC3 methods, SIR classification was determined using NARMS breakpoints for *Salmonella* (accessed March 23, 2020).

### 3.3 Genomic AMR determinants of bovine-associated *Salmonella* are serotype-associated

Based on the presence and absence of pan-genome elements among all 128 *Salmonella* isolates sequenced here, the *Salmonella* pan-genome was more similar within serotype and within farm than between serotype and between farm, respectively (PERMANOVA and ANOSIM *P* < 0.05 after a Bonferroni correction; Figure 4 and Table 2), with serotypes showing a higher degree of pan-genome dissimilarity (ANOSIM *R* = 0.99) and accounting for a larger proportion of the variance (PERMANOVA *R^2^* = 0.93) than farms (Figure 4 and Table 2); however, dispersion among both serotypes and farms differed (PERMDISP2 *P* < 0.05 after a Bonferrroni correction; Table 2), indicating that the ANOSIM and/or PERMANOVA tests could potentially be confounding dispersion with serotype/farm. Additionally, subclinical bovine *Salmonella* isolates did not significantly differ from strains isolated from the associated farm environment based on pan-genome element presence/absence (PERMANOVA, ANOSIM, and PERMDISP2 *P* > 0.05 after a Bonferroni correction; Figure 4 and Table 2).

**Figure 4.**
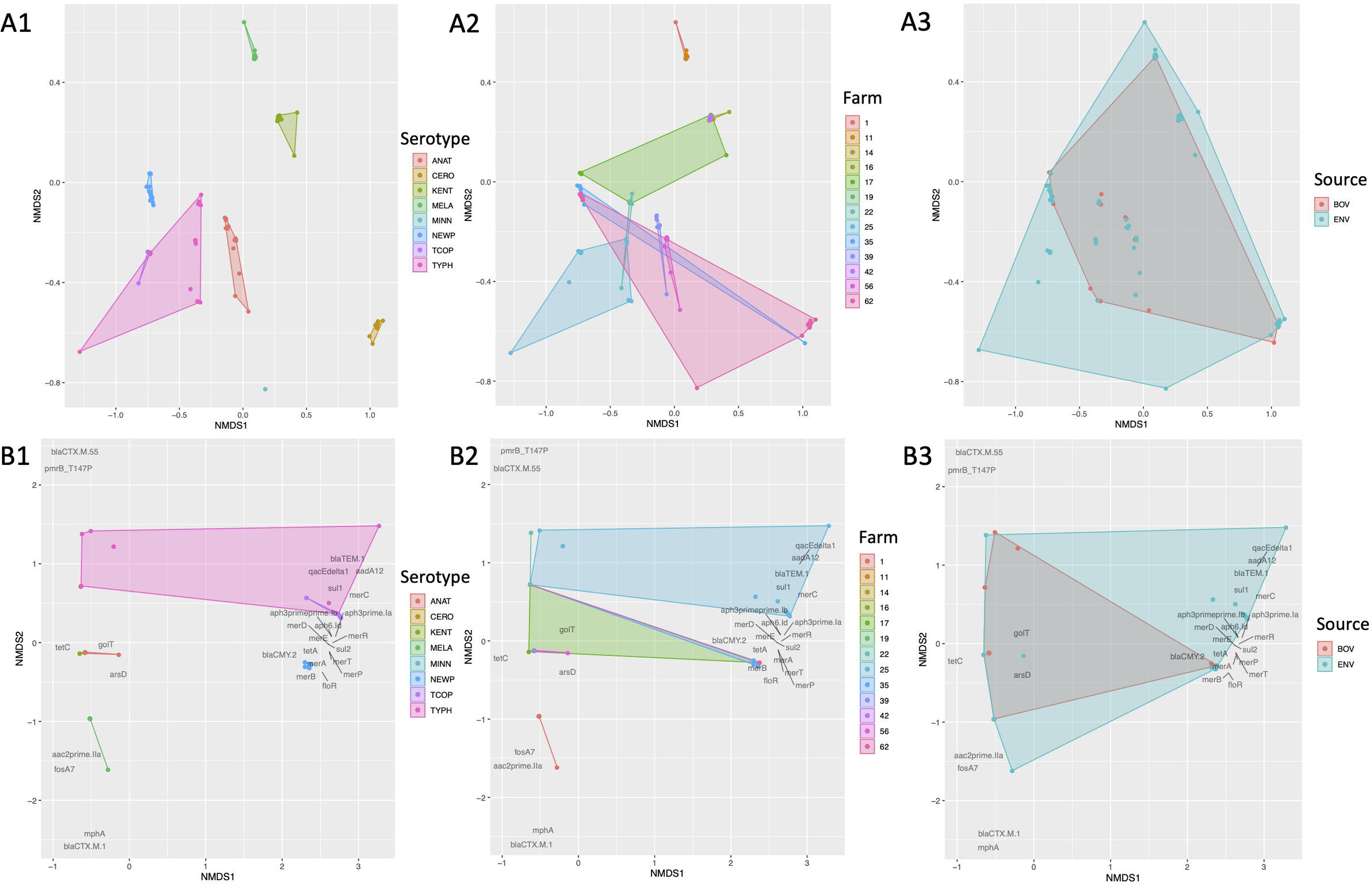
Results of nonmetric multidimensional scaling (NMDS) performed using the presence and absence of (A) pan-genome elements (*n* = 4,102; identified using Roary), and (B) antimicrobial resistance (AMR) and stress response genes (*n* = 28; detected using AMRFinderPlus) among 128 bovine-associated *Salmonella* isolates, plotted in two dimensions. Points represent isolates, while shaded regions and convex hulls correspond to isolate (1) serotypes (ANAT, Anatum; CERO, Cerro; KENT, Kentucky; MELA, Meleagridis; NEWP, Newport; TCOP, Typhimurium Copenhagen; TYPH, Typhimurium), (2) farm, and (3) source (BOV, bovine; ENV, bovine farm environment). For all plots, a Jaccard distance metric was used. For AMR/stress response genes (B), gene names/scores are plotted in dark gray text.

**Table 2.** Results of PERMDISP2, ANOSIM, and PERMANOVA tests.^a^

| Group | PERMDISP2 Raw<br>*P*-Value (*F*)<sup>b</sup> | ANOSIM Raw *P*-<br>Value (*R*)<sup>c</sup> | PERMANOVA Raw *P*-<br>Value (*R*<sup>2</sup>)<sup>d</sup> |
| --- | --- | --- | --- |
| Pan-genome element presence/absence (*n* = 4,102)<sup>e</sup> |  |  |  |
| Serotype | 2.0E-4 (14.3)* | < 1.0E-4 (0.99)* | < 1.0E-4 (0.93)* |
| Farm | < 1.0E-4 (4.43)* | < 1.0E-4 (0.54)* | < 1.0E-4 (0.73)* |
| Source | 0.071 (3.40) | 0.99 (-0.07) | 0.32 (0.01) |
| Antimicrobial resistance and stress response gene presence/absence (*n* = 28)<sup>f</sup> |  |  |  |
| Serotype | 0.013 (4.46) | < 1.0E-4 (0.79)* | < 1.0E-4 (0.85)* |
| Farm | < 1.0E-4 (5.52)* | < 1.0E-4 (0.30)* | < 1.0E-4 (0.54)* |
| Source | 0.74 (0.01) | 0.79 (-0.03) | 0.31 (0.01) |

<sup>a</sup>All tests were performed using a Jaccard dissimilarity metric and 10,000 permutations; raw *P*-values are reported for all tests, with significant *P*-values (*P* < 0.05 after a Bonferroni correction was applied to all values) denoted with an asterisk (\*).
<sup>b</sup>ANOVA-like permutation test applied to results obtained using the PERMDISP2 procedure for the analysis of multivariate homogeneity of group dispersions (i.e., variances), obtained using the betadisper and permutest functions in the vegan package in R; betadisper is a multivariate analogue of Levene's test for homogeneity of variances.
<sup>c</sup>Analysis of similarities (ANOSIM) test results obtained using the anosim function in the vegan package in R
<sup>d</sup>Permutational analysis of variance (PERMANOVA) test results obtained using the adonis2 function in the vegan package in R
<sup>e</sup>Identified using Roary and a 70% protein BLAST (BLASTP) identity threshold
<sup>f</sup>Detected using AMRFinderPlus

Based on the presence and absence of AMR and stress response determinants detected among all 128 *Salmonella* genomes, isolates were more similar within serotype than between serotypes (PERMANOVA and ANOSIM *P* < 0.05 and PERMDISP2 *P* > 0.05 after a Bonferroni correction; Figure 4 and Table 2). Additionally, isolates were more similar within farm than between farm based on their AMR and stress response gene presence/absence profiles (PERMANOVA and ANOSIM *P* < 0.05; Figure 4 and Table 2), although significant, potentially confounding dispersion differences among farms were present (PERMDISP2 *P* < 0.05; Table 2). As was the case with the pan-genome in its entirety, subclinical bovine *Salmonella* isolates did not significantly differ from farm environmental isolates based on AMR and stress response gene presence/absence (PERMANOVA, ANOSIM, and PERMDISP2 *P* > 0.05 after a Bonferroni correction; Figure 4 and Table 2).

### 3.4 Each of two New York State dairy farms harbors a unique, bovine-associated *S.* Anatum lineage

Fifteen *S.* Anatum strains encompassing four PFGE types (Supplementary Table S1) were isolated from subclinical bovine sources and their associated farm environments on two different New York State dairy farms (i.e., Farms 39 and 56; Figure 5 and Table 3). Notably, the *S.* Anatum lineages circulating on each farm were distinct at a genomic level, with isolates from each farm forming a separate clade (posterior probability [PP] = 1 for each; Figure 5). The two farm-associated lineages were predicted to share a common ancestor circa 1836 (node age 1836.28 using median node heights; Figure 5), although the age of the common ancestor could not be dated reliably (node height 95% highest posterior density [HPD] interval 540.85-1978.42; Supplementary Figure S5).

**Figure 5.**
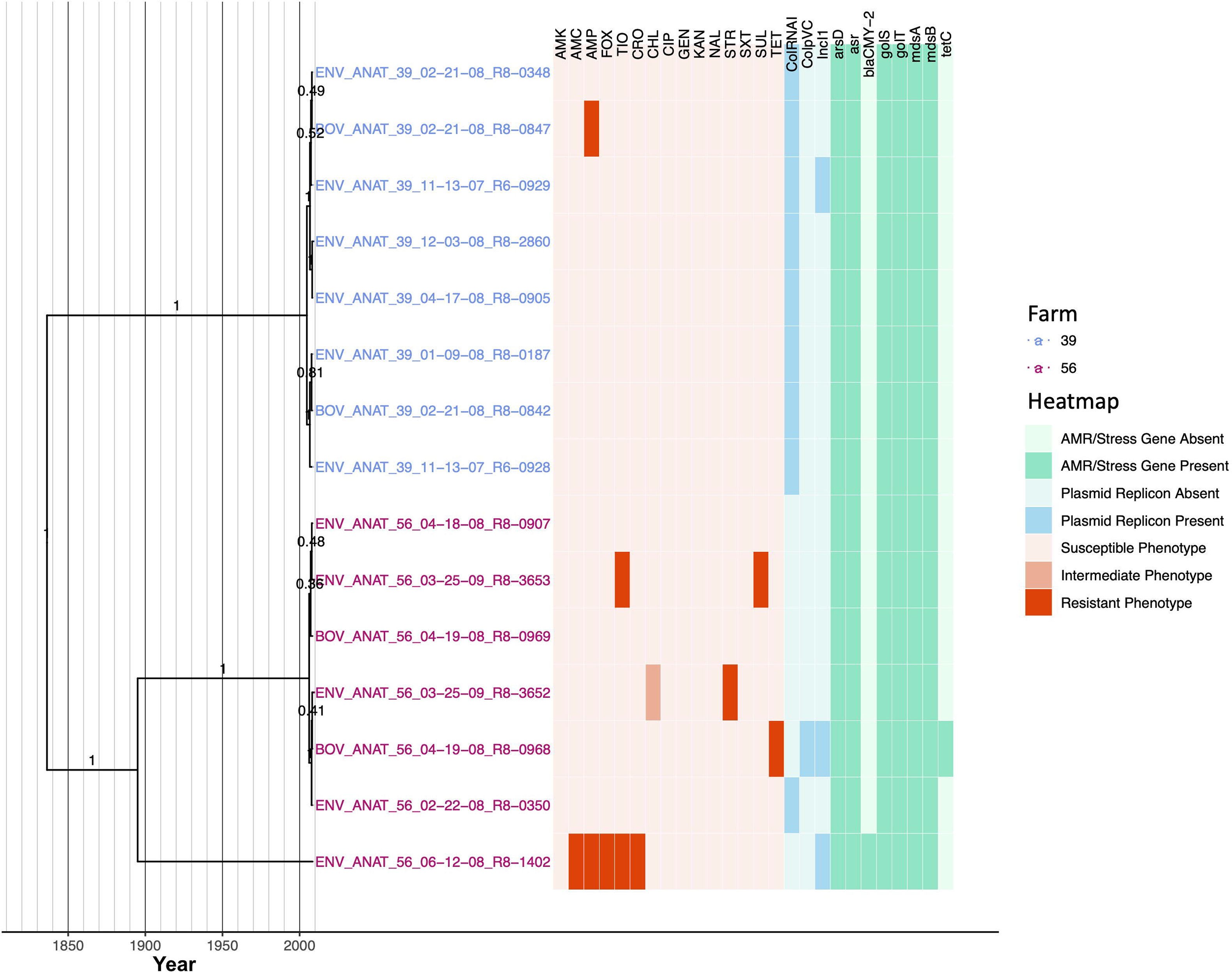
Rooted, time-scaled maximum clade credibility (MCC) phylogeny constructed using core SNPs identified among 15 *Salmonella* Anatum genomes isolated from subclinical bovine sources and the surrounding bovine farm environment. Tip label colors denote the ID of the farm from which each strain was isolated. Branch labels denote posterior probabilities of branch support. Time in years is plotted along the X-axis, and branch lengths are reported in years. The heatmap to the right of the phylogeny denotes (i) the susceptible-intermediate-resistant (SIR) classification of each isolate for each of 15 antimicrobials (obtained using phenotypic testing and NARMS breakpoints; orange); (ii) presence and absence of plasmid replicons (detected using ABRicate/PlasmidFinder and minimum nucleotide identity and coverage thresholds of 80 and 60%, respectively; blue); (iii) presence and absence of antimicrobial resistance (AMR) and stress response genes (identified using AMRFinderPlus and default parameters; green). Core SNPs were identified using Snippy. The phylogeny was constructed using the results of ten independent runs using a strict clock model, the Standard_TVMef nucleotide substitution model, and the Coalescent Bayesian Skyline population model implemented in BEAST version 2.5.1, with 10% burn-in applied to each run. LogCombiner-2 was used to combine BEAST 2 log files, and TreeAnnotator-2 was used to construct the phylogeny using median node heights. Abbreviations for the 15 antimicrobials are: AMK, amikacin; AMC, amoxicillin-clavulanic acid; AMP, ampicillin; FOX, cefoxitin; TIO, ceftiofur; CRO, ceftriaxone; CHL, chloramphenicol; CIP, ciprofloxacin; GEN, gentamicin; KAN, kanamycin; NAL, nalidixic acid; STR, streptomycin; SXT, sulfamethoxazole-trimethoprim; SUL, sulfisoxazole; TET, tetracycline.

**Table 3.** Summary of within-serotype evolutionary analyses.^a^

| Serotype | Isolates | Core SNPs (Pairwise Range)<sup>b</sup> | Clock/Population Model<sup>c</sup> | Mean/Median Tree Height in Years (95% HPD Interval)<sup>d</sup> | Mean/Median Evolutionary Rate in Substitutions/Site/Year (95% HPD Interval)<sup>e</sup> |
| --- | --- | --- | --- | --- | --- |
| Anatum | 15 | 337 (0-257) | Strict/Skyline | 1484.9/1837.0 (549.6-1980.1) | $1.67 \times 10^{-7} / 1.48 \times 10^{-7}$ ($6.92 \times 10^{-11} - 3.86 \times 10^{-7}$) |
| Cerro | 13 | 21 (0-12) | Strict/Skyline | 2008.2/2008.4 (2007.6-2008.6) | $9.11 \times 10^{-7} / 8.94 \times 10^{-7}$ ($3.06 \times 10^{-7} - 1.57 \times 10^{-6}$) |
| Kentucky | 36 | 102 (0-30) | Relaxed/Skyline | 2004.1/2005.0 (2000.8-2006.8) | $6.39 \times 10^{-7} / 6.34 \times 10^{-7}$ ($2.05 \times 10^{-7} - 1.07 \times 10^{-6}$) |
| Meleagridis | 19 | 27 (0-9) | Strict/Skyline | 2007.4/2007.5 (2006.9-2007.7) | $6.88 \times 10^{-7} / 6.66 \times 10^{-7}$ ($2.90 \times 10^{-7} - 1.12 \times 10^{-6}$) |
| Newport | 16 | 52 (0-38) | Relaxed/Skyline | 2004.2/2004.6 (2000.4-2007.8) | $9.02 \times 10^{-7} / 8.22 \times 10^{-7}$ ($2.64 \times 10^{-7} - 1.65 \times 10^{-6}$) |
| Typhimurium (Copenhagen) | 27 | 732 (0-634) | Relaxed/Skyline | 1936.0/1943.0 (1864.7-1991.4) | $1.07 \times 10^{-6} / 9.66 \times 10^{-7}$ ($2.84 \times 10^{-7} - 2.05 \times 10^{-6}$) |

<sup>a</sup>See Supplementary Table S5 for an extended version of this table; note that evolutionary rates may be higher than previously reported estimates for *Salmonella* populations isolated over a longer time frame, due to the small sample sizes and short temporal period characterized here (Moller et al., 2018).
<sup>b</sup>Number of core SNPs identified among all genomes within the serotype after removing recombination with Gubbins; the range of pairwise SNP differences between isolates was calculated using the dist.gene function in the ape package in R.
<sup>c</sup>The optimal model selected for the data set; can be a combination of a strict or lognormal relaxed molecular clock (“Strict” or “Relaxed”, respectively) and a Constant Coalescent or Coalescent Bayesian Skyline population model (“CC” or “Skyline”, respectively); see Supplementary Table S5 for more details.
<sup>d</sup>The tree height parameter and its respective 95% highest posterior density (HPD) interval reported by Tracer.
<sup>e</sup>Corresponds to the clockRate and rate.mean parameters estimated by BEAST2 for models using strict and lognormal relaxed molecular clock models, respectively, as reported by Tracer.

*S.* Anatum isolates from Farm 39 shared a common ancestor circa 2005 (node age 2004.69, node height 95% HPD 1978.46-2007.69; Figure 5 and Supplementary Figure S5). All Farm 39 *S.* Anatum isolates possessed identical AMR/stress response gene profiles, and all isolates were pan-susceptible except for a single isolate that was resistant to ampicillin (Figure 5). All Farm 39 *S.* Anatum isolates additionally harbored ColRNAI plasmids; a single isolate additionally harbored an IncI1 plasmid that appeared to harbor no AMR genes (Figure 5).

*S.* Anatum isolates from Farm 56, however, were considerably more diverse than their Farm 39 counterparts; while a clade containing six of seven strains shared a very recent common ancestor (node age 2006.16, node height 95% HPD 1985.37-2008.11; Figure 5 and Supplementary Figure S5), a unique lineage represented by a single environmental isolate (ENV_ANAT_56_06-12-08_R8-1402) was present among *S.* Anatum from Farm 56 (Figure 5). All *S.* Anatum isolates from Farm 56 were predicted to have evolved from a common ancestor that existed circa 1895 (node age 1895.03), although this node could not be reliably dated (node height 95% HPD 1036.89-1989.944; Supplementary Figure S5). Additionally, *S.* Anatum isolated from Farm 56 showcased a greater degree of AMR heterogeneity than those from Farm 39 (Figure 5). Notably, the isolate comprising the unique Farm 56 *S.* Anatum lineage possessed an IncI1 plasmid and *bla*_CMY-2_ and was multidrug resistant (MDR; resistant to amoxicillin-clavulanic acid, ampicillin, cefoxitin, ceftiofur, and ceftriaxone; Figure 5). Three of six *S.* Anatum strains comprising the major Farm 56 *S.* Anatum lineage were pan-susceptible. The remaining three isolates were resistant to one of (i) tetracycline, (ii) streptomycin, or (iii) ceftiofur and sulfisoxazole; the streptomycin-resistant isolate additionally exhibited reduced susceptibility to chloramphenicol (Figure 5). The tetracycline-resistant isolate additionally possessed both ColpVC and IncI1plasmids and harbored tetracycline resistance gene *tetC* (Figure 5). While these data suggest some *S.* Anatum lineages queried here have recently acquired AMR, the limited number of isolates and the large degree of uncertainty for some phylogeny node ages preclude reliable estimation of AMR acquisition timeframes.

### 3.5 A closely related *S.* Cerro lineage spans two New York State dairy farms

Thirteen *S.* Cerro strains encompassing two PFGE types (Supplementary Table S1) isolated from two dairy farms (12 from Farm 62 and one from Farm 35) were found to share a high degree of genomic similarity; isolates differed by, at most, 12 core SNPs and evolved from a common ancestor that existed circa March 2008 (node age 2008.21, common ancestor [CA] node height 95% HPD interval 2007.6-2008.6; Figure 6, Table 3, Supplementary Figure S6, and Supplementary Table S5). While IncI1 and ColRNAI plasmid replicons were detected in all 13 *S.* Cerro isolates, only one isolate was not pan-susceptible (Figure 6). Notably, the isolate from Farm 35 (BOV_CERO_35_10-02-08_R8-2685) was classified as resistant to nine antimicrobials using phenotypic methods (i.e., amoxicillin-clavulanic acid, ampicillin, cefoxitin, ceftiofur, ceftriaxone, chloramphenicol, streptomycin, sulfisoxazole, and tetracycline); based on the most parsimonious explanation for AMR acquisition, this lineage acquired AMR after July 2008 (node age 2008.51, CA node height 95% HPD interval 2008.14-2008.75; Figure 6 and Supplementary Figure S6). However, no genomic determinants known to confer resistance to these antimicrobials were detected in the genome of the MDR isolate (Figure 6), and the MDR phenotype was confirmed in a second, independent phenotypic AMR test (Supplementary Table S4).

**Figure 6.**
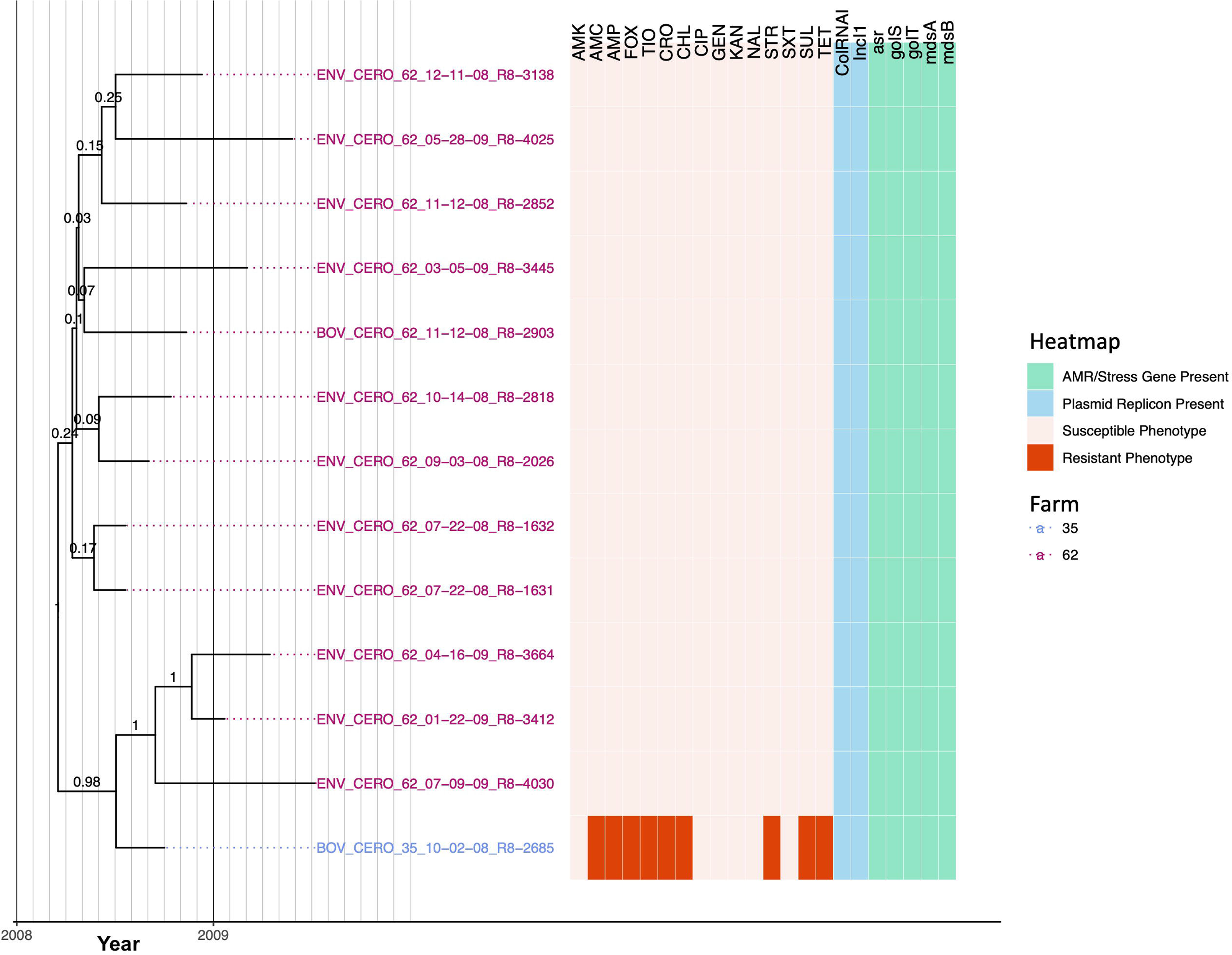
Rooted, time-scaled maximum clade credibility (MCC) phylogeny constructed using core SNPs identified among 13 *Salmonella* Cerro genomes isolated from subclinical bovine sources and the surrounding bovine farm environment. Tip label colors denote the ID of the farm from which each strain was isolated. Branch labels denote posterior probabilities of branch support. Time in years is plotted along the X-axis, and branch lengths are reported in years. The heatmap to the right of the phylogeny denotes (i) the susceptible-intermediate-resistant (SIR) classification of each isolate for each of 15 antimicrobials (obtained using phenotypic testing and NARMS breakpoints; orange); (ii) presence of plasmid replicons (detected using ABRicate/PlasmidFinder and minimum nucleotide identity and coverage thresholds of 80 and 60%, respectively; blue); (iii) presence of antimicrobial resistance (AMR) and stress response genes (identified using AMRFinderPlus and default parameters; green). Core SNPs were identified using Snippy. The phylogeny was constructed using the results of ten independent runs using a strict clock model, the Standard_TPM1 nucleotide substitution model, and the Coalescent Bayesian Skyline population model implemented in BEAST version 2.5.1, with 10% burn-in applied to each run. LogCombiner-2 was used to combine BEAST 2 log files, and TreeAnnotator-2 was used to construct the phylogeny using common ancestor node heights. Abbreviations for the 15 antimicrobials are: AMK, amikacin; AMC, amoxicillin-clavulanic acid; AMP, ampicillin; FOX, cefoxitin; TIO, ceftiofur; CRO, ceftriaxone; CHL, chloramphenicol; CIP, ciprofloxacin; GEN, gentamicin; KAN, kanamycin; NAL, nalidixic acid; STR, streptomycin; SXT, sulfamethoxazole-trimethoprim; SUL, sulfisoxazole; TET, tetracycline.

### 3.6 *S.* Kentucky strains isolated across five different New York State dairy farms evolved from a common ancestor that existed circa 2004

Thirty-six *S.* Kentucky isolates encompassing two PFGE types (Supplementary Table S1) isolated across five New York State dairy farms (i.e., five, seven, nine, seven, and eight isolates from each of Farm 14, 16, 17, 19, and 42, respectively) were similar at a genomic level; isolates differed by between 0 and 30 core SNPs and shared a common ancestor that was predicted to have existed circa January/February 2004 (node age 2004.07, CA node height 95% HPD interval 2000.73-2006.8; Figure 7, Table 3, Supplementary Figure S7, and Supplementary Table S5). Two farms harbored a total of three *S.* Kentucky isolates, which were not pan-susceptible (two isolates from Farm 16 and one from Farm 17; Figure 7). Farm 17 harbored a tetracycline-resistant isolate (ENV_KENT_17_03-11-08_R8-0815), which possessed an IncI1 plasmid and *tetC* (Figure 7). The lineage represented by this isolate was predicted to have acquired tetracycline resistance after March 2007 (node height 2007.19, CA node height 95% HPD interval 2006.43-2007.84; Figure 7 and Supplementary Figure S7). The two *S.* Kentucky isolates from Farm 16 additionally showed reduced susceptibility to chloramphenicol, a trait predicted to have been acquired by these lineages after December 2006/January 2007 (for the lineage represented by isolate ENV_KENT_16_12-04-07_R8-0061; node height 2006.98, CA node height 95% HPD interval 2005.95-2007.85) and May 2007 (for the lineage represented by isolate BOV_KENT_16_02-13-08_R8-0838; node height 2007.38, CA node height 95% HPD interval 2006.59-2008.10; Figure 7 and Supplementary Figure S7). No corresponding genes that may encode for reduced chloramphenicol susceptibility were identified in these two isolates.

**Figure 7.**
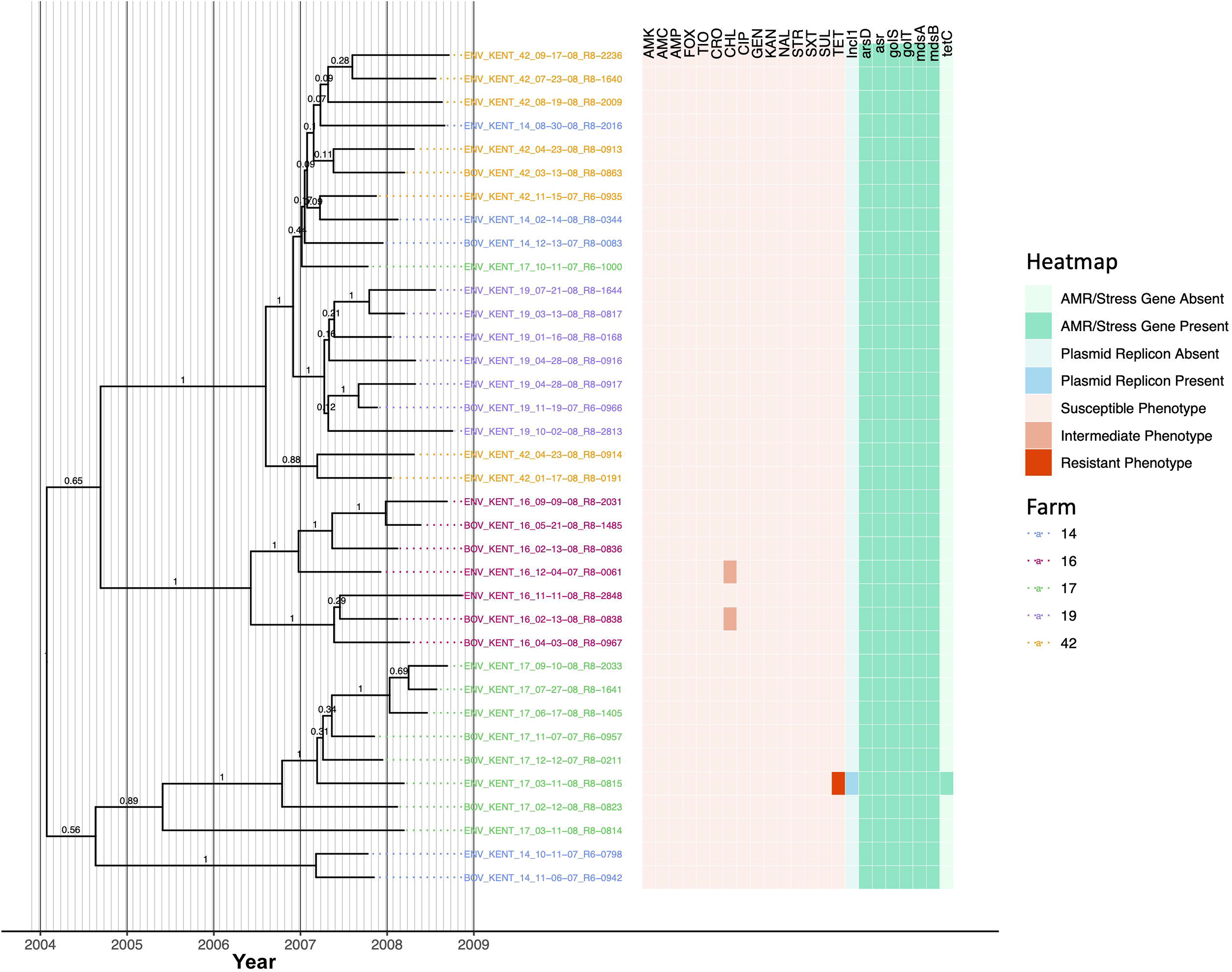
Rooted, time-scaled maximum clade credibility (MCC) phylogeny constructed using core SNPs identified among 36 *Salmonella* Kentucky genomes isolated from subclinical bovine sources and the surrounding bovine farm environment. Tip label colors denote the ID of the farm from which each strain was isolated. Branch labels denote posterior probabilities of branch support. Time in years is plotted along the X-axis, and branch lengths are reported in years. The heatmap to the right of the phylogeny denotes (i) the susceptible-intermediate-resistant (SIR) classification of each isolate for each of 15 antimicrobials (obtained using phenotypic testing and NARMS breakpoints; orange); (ii) presence of plasmid replicons (detected using ABRicate/PlasmidFinder and minimum nucleotide identity and coverage thresholds of 80 and 60%, respectively; blue); (iii) presence of antimicrobial resistance (AMR) and stress response genes (identified using AMRFinderPlus and default parameters; green). Core SNPs were identified using Snippy. The phylogeny was constructed using the results of ten independent runs using a relaxed lognormal clock model, the Standard_TVMef nucleotide substitution model, and the Coalescent Bayesian Skyline population model implemented in BEAST version 2.5.1, with 10% burn-in applied to each run. LogCombiner-2 was used to combine BEAST 2 log files, and TreeAnnotator-2 was used to construct the phylogeny using common ancestor node heights. Abbreviations for the 15 antimicrobials are: AMK, amikacin; AMC, amoxicillin-clavulanic acid; AMP, ampicillin; FOX, cefoxitin; TIO, ceftiofur; CRO, ceftriaxone; CHL, chloramphenicol; CIP, ciprofloxacin; GEN, gentamicin; KAN, kanamycin; NAL, nalidixic acid; STR, streptomycin; SXT, sulfamethoxazole-trimethoprim; SUL, sulfisoxazole; TET, tetracycline.

### 3.7 A clonal *S.* Meleagridis lineage is distributed across two New York State dairy farms and encompasses isolates carrying *bla*_CTX-M-1_

Nineteen *S.* Meleagridis isolates encompassing two PFGE types (Supplementary Table S1) were isolated from two dairy farms (13 and six isolates from Farms 01 and 11, respectively) and were highly clonal: isolates differed by fewer than ten core SNPs and evolved from a common ancestor that existed circa May/June 2007 (node age 2007.42, CA node height 95% HPD interval 2006.91-2007.75; Figure 8, Table 3, Supplementary Figure S8, and Supplementary Table S5). All but three branches within the *S.* Meleagridis phylogeny had low support (PP ≤ 0.41; Figure 8 and Supplementary Figure S8), indicating that most nodes were unreliable, likely due to the isolates being highly clonal. All *S.* Meleagridis isolates from Farm 11 were pan-susceptible, possessed no plasmid replicons, and did not possess any acquired AMR genes (Figure 8). Among the *S.* Meleagridis isolates from Farm 01, one isolate (ENV_MELA_01_10-02-07_R6-0938) was resistant to ampicillin, ceftiofur, and ceftriaxone, and possessed an IncN plasmid, macrolide resistance gene *mph(A)*, and beta-lactamase *bla*_CTX-M-1_ (Figure 8). Two additional *S.* Meleagridis isolates from Farm 01 each exhibited reduced susceptibility to either (i) cefoxitin, sulfisoxazole, and tetracycline, or (ii) ceftiofur (Figure 8).

**Figure 8.**
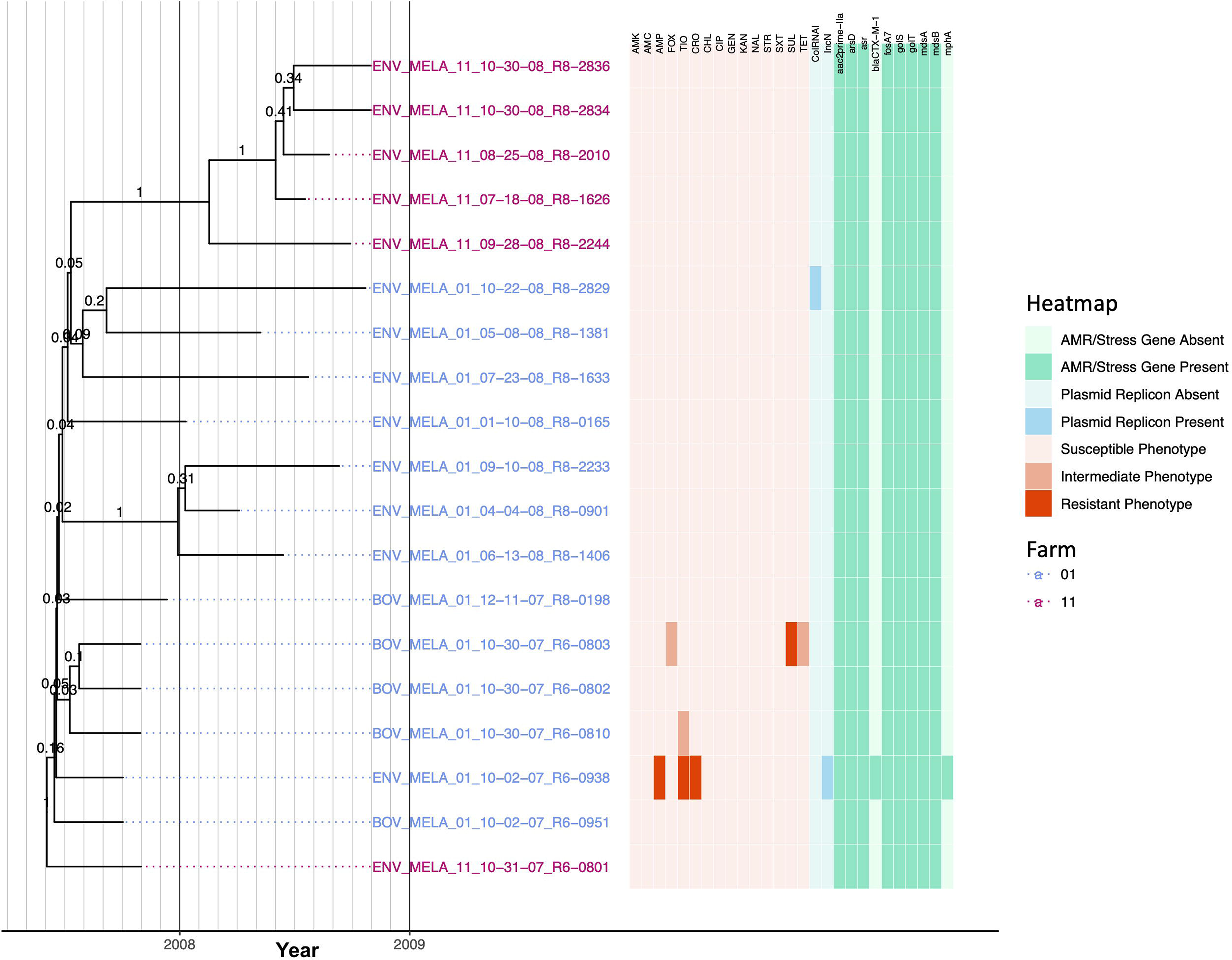
Rooted, time-scaled maximum clade credibility (MCC) phylogeny constructed using core SNPs identified among 19 *Salmonella* Meleagridis genomes isolated from subclinical bovine sources and the surrounding bovine farm environment. Tip label colors denote the ID of the farm from which each strain was isolated. Branch labels denote posterior probabilities of branch support. Time in years is plotted along the X-axis, and branch lengths are reported in years. The heatmap to the right of the phylogeny denotes (i) the susceptible-intermediate-resistant (SIR) classification of each isolate for each of 15 antimicrobials (obtained using phenotypic testing and NARMS breakpoints; orange); (ii) presence and absence of plasmid replicons (detected using ABRicate/PlasmidFinder and minimum nucleotide identity and coverage thresholds of 80 and 60%, respectively; blue); (iii) presence and absence of antimicrobial resistance (AMR) and stress response genes (identified using AMRFinderPlus and default parameters; green). Core SNPs were identified using Snippy. The phylogeny was constructed using the results of ten independent runs using a strict clock model, the Standard_TPM2 nucleotide substitution model, and the Coalescent Bayesian Skyline population model implemented in BEAST version 2.5.1, with 10% burn-in applied to each run. LogCombiner-2 was used to combine BEAST 2 log files, and TreeAnnotator-2 was used to construct the phylogeny using common ancestor node heights. Abbreviations for the 15 antimicrobials are: AMK, amikacin; AMC, amoxicillin-clavulanic acid; AMP, ampicillin; FOX, cefoxitin; TIO, ceftiofur; CRO, ceftriaxone; CHL, chloramphenicol; CIP, ciprofloxacin; GEN, gentamicin; KAN, kanamycin; NAL, nalidixic acid; STR, streptomycin; SXT, sulfamethoxazole-trimethoprim; SUL, sulfisoxazole; TET, tetracycline.

### 3.8 Kanamycin resistance among each of three New York State dairy farms harboring a distinct, multidrug-resistant *S.* Newport lineage is farm-associated

Sixteen *S.* Newport isolates encompassing three PFGE types (Supplementary Table S1) were isolated from one of three farms (four, five, and seven isolates from Farms 17, 35, and 62, respectively); all isolates were resistant to amoxicillin-clavulanic acid, ampicillin, cefoxitin, ceftiofur, ceftriaxone, streptomycin, sulfisoxazole, and tetracycline (Figure 9). All *S.* Newport genomes harbored IncA/C2 and ColRNAI plasmids, as well as streptomycin resistance genes *APH(3’’)-Ib* and *APH(6)-Id* (i.e., *strAB*), beta-lactamase *bla*_CMY-2_, sulfonamide resistance gene *sul2*, and tetracycline resistance gene *tetA* (Figure 9). Notably, the *S.* Newport lineage circulating on each farm formed one of three separate clades (PP = 0.99-1.0) that evolved from a common ancestor that existed circa March/April 2004 (node age 2004.23, CA node height 95% HPD interval 2000.42-2007.85; Figure 9, Table 3, Supplementary Figure S9, and Supplementary Table S5).

**Figure 9.**
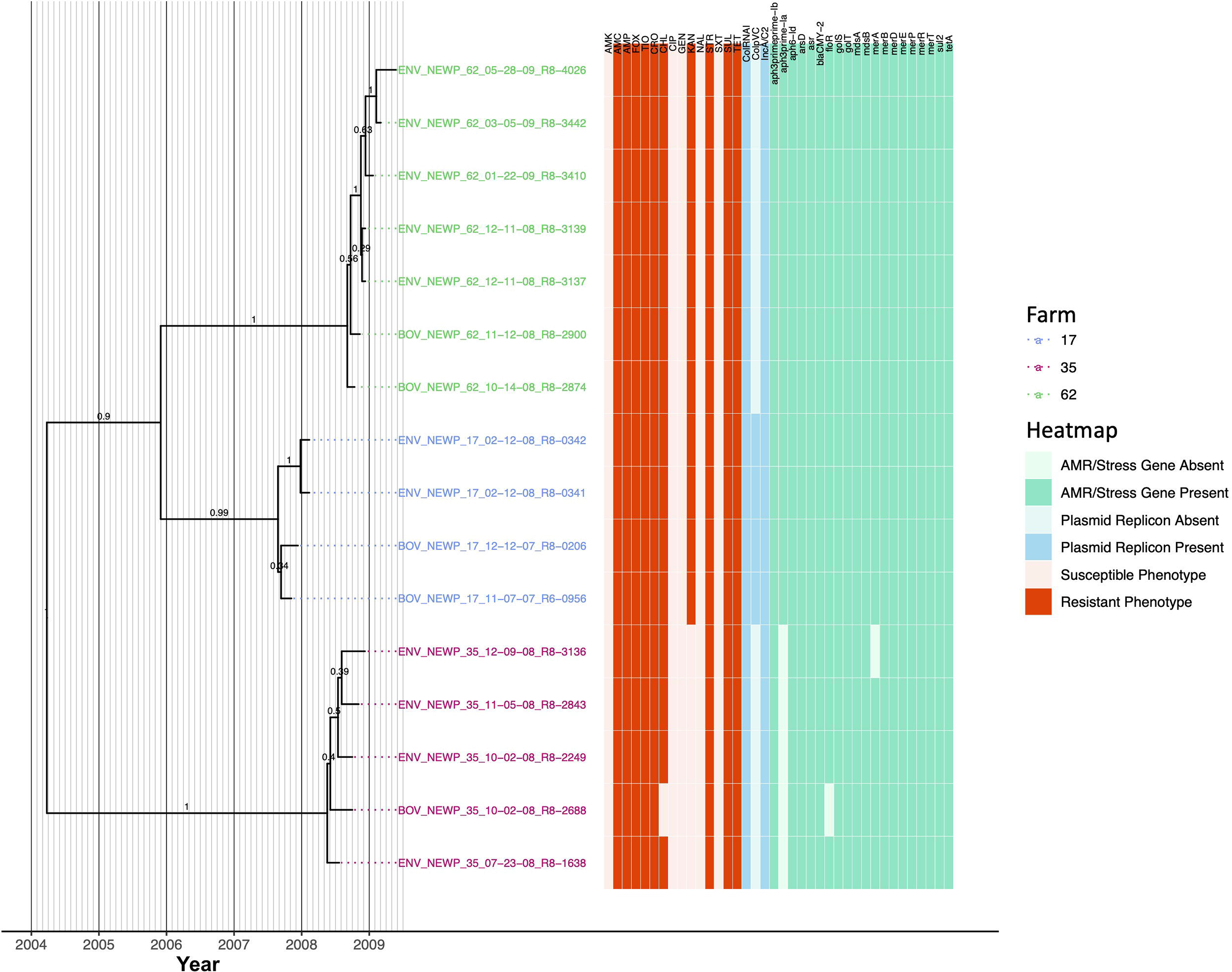
Rooted, time-scaled maximum clade credibility (MCC) phylogeny constructed using core SNPs identified among 16 *Salmonella* Newport genomes isolated from subclinical bovine sources and the surrounding bovine farm environment. Tip label colors denote the ID of the farm from which each strain was isolated. Branch labels denote posterior probabilities of branch support. Time in years is plotted along the X-axis, and branch lengths are reported in years. The heatmap to the right of the phylogeny denotes (i) the susceptible-intermediate-resistant (SIR) classification of each isolate for each of 15 antimicrobials (obtained using phenotypic testing and NARMS breakpoints; orange); (ii) presence and absence of plasmid replicons (detected using ABRicate/PlasmidFinder and minimum nucleotide identity and coverage thresholds of 80 and 60%, respectively; blue); (iii) presence and absence of antimicrobial resistance (AMR) and stress response genes (identified using AMRFinderPlus and default parameters; green). Core SNPs were identified using Snippy. The phylogeny was constructed using the results of ten independent runs using a relaxed lognormal clock model, the Standard_TPM1 nucleotide substitution model, and the Coalescent Bayesian Skyline population model implemented in BEAST version 2.5.1, with 10% burn-in applied to each run. LogCombiner-2 was used to combine BEAST 2 log files, and TreeAnnotator-2 was used to construct the phylogeny using common ancestor node heights. Abbreviations for the 15 antimicrobials are: AMK, amikacin; AMC, amoxicillin-clavulanic acid; AMP, ampicillin; FOX, cefoxitin; TIO, ceftiofur; CRO, ceftriaxone; CHL, chloramphenicol; CIP, ciprofloxacin; GEN, gentamicin; KAN, kanamycin; NAL, nalidixic acid; STR, streptomycin; SXT, sulfamethoxazole-trimethoprim; SUL, sulfisoxazole; TET, tetracycline.

The *S.* Newport lineages present on Farm 17 and Farm 62 were additionally resistant to chloramphenicol and kanamycin and possessed chloramphenicol and kanamycin resistance genes *floR* and *APH(3’)-Ia*, respectively (Figure 9). The Farm 17 and Farm 62 lineages evolved from a common ancestor predicted to have existed circa November/December 2005 (node age 2005.91, CA node height 95% HPD interval 2003.77-2007.85; Figure 9 and Supplementary Figure S9). All members of the Farm 17 lineage additionally harbored a ColpVC plasmid and shared a common ancestor dated to circa August/September 2007 (node age 2007.65, CA node height 95% HPD interval 2007.29-2007.85; Figure 9 and Supplementary Figure S9). The Farm 62 lineage, which did not possess the ColpVC plasmid, evolved from a common ancestor circa August/September 2008 (node age 2008.68, CA node height 95% HPD interval 2008.42-2008.78; Figure 9 and Supplementary Figure S9).

Unlike the *S.* Newport lineages present on Farm 17 and Farm 62, the Farm 35 *S.* Newport lineage did not possess kanamycin resistance gene *APH(3’)-Ia* and was kanamycin-susceptible (Figure 9). The common ancestor of the Farm 35 *S.* Newport lineage was dated circa May 2008 (node age 2008.38, CA node height 95% HPD interval 2008.06-2008.55). All but one Farm 35 *S.* Newport isolates were additionally resistant to chloramphenicol and possessed *floR*; BOV_NEWP_35_10-02-08_R8-2688 did not possess *floR* and was chloramphenicol-susceptible (Figure 9).

### 3.9 Each of four major lineages composed of *S.* Typhimurium and its O5-Copenhagen variant is associated with one of three New York State dairy farms

Twenty-seven bovine and farm environmental *S.* Typhimurium and *S.* Typhimurium Copenhagen isolates that encompassed five PFGE types (Supplementary Table S1) were isolated from three dairy farms (1, 10, and 16 strains isolated from Farm 17, 22, and 25, respectively). All isolates queried here shared a common ancestor that existed circa 1936 (node age 1935.62, CA node height 95% HPD interval 1864.84-1991.86; Figure 10, Table 3, Supplementary Figure S10, and Supplementary Table S5). Notably, the *S.* Typhimurium Copenhagen variant was polyphyletic (Figure 10), regardless of whether traditional or *in silico* (i.e., SeqSero2) methods had been used for serotype variant assignment. Additionally, the *S.* Typhimurium/*S.* Typhimurium Copenhagen isolates sequenced here showcased the most diverse AMR phenotypic profiles and AMR gene presence/absence profiles (Figures 4 and 10).

**Figure 10.**
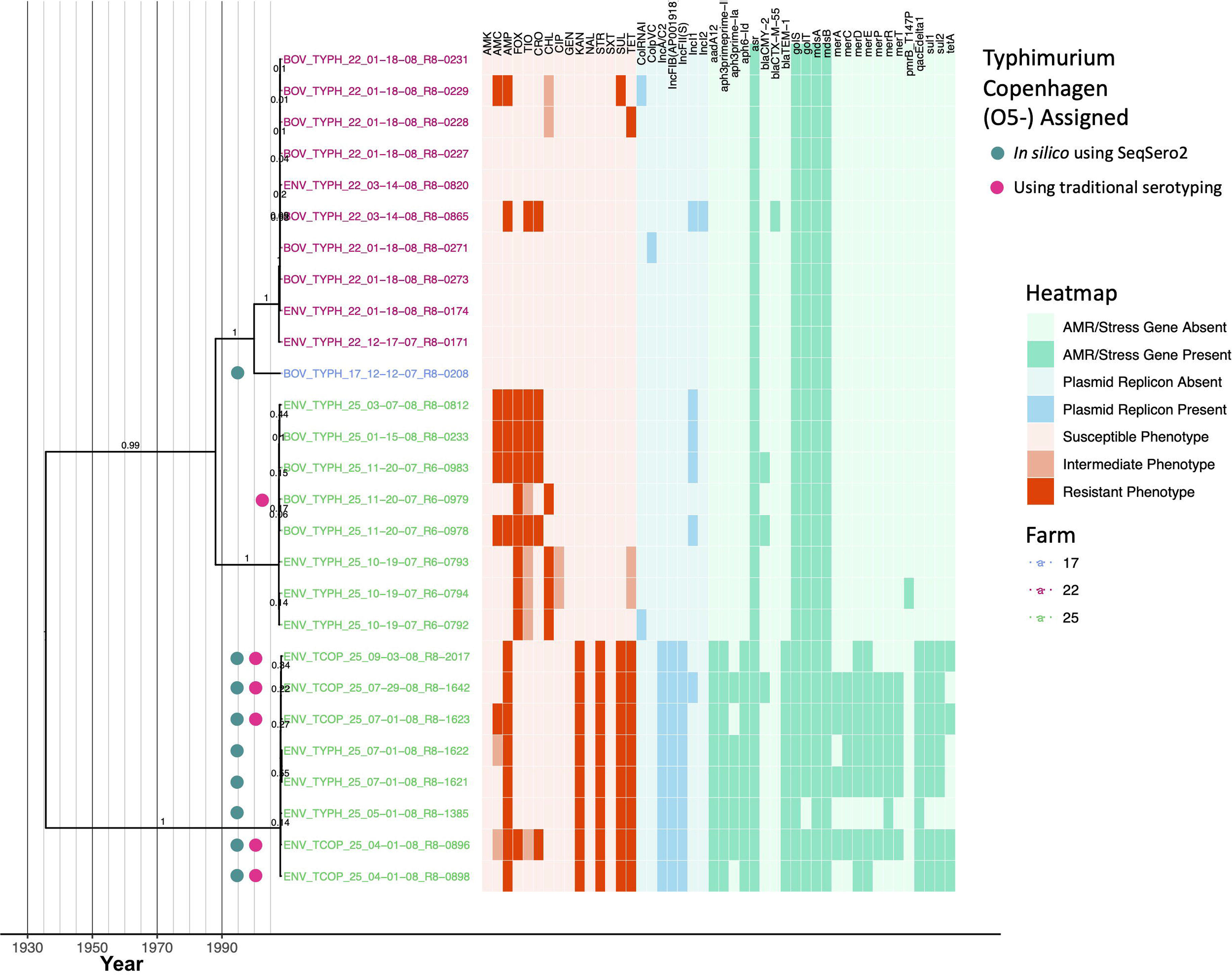
Rooted, time-scaled maximum clade credibility (MCC) phylogeny constructed using core SNPs identified among 27 *Salmonella* Typhimurium and Typhimurium Copenhagen genomes isolated from subclinical bovine sources and the surrounding bovine farm environment. Tip label colors denote the ID of the farm from which each strain was isolated. Circles to the left of tip labels denote isolates that were assigned to the Typhimurium Copenhagen variant of *S.* Typhimurium using SeqSero2 (teal) and/or traditional serotyping (pink). Branch labels denote posterior probabilities of branch support. Time in years is plotted along the X-axis, and branch lengths are reported in years. The heatmap to the right of the phylogeny denotes (i) the susceptible-intermediate-resistant (SIR) classification of each isolate for each of 15 antimicrobials (obtained using phenotypic testing and NARMS breakpoints; orange); (ii) presence and absence of plasmid replicons (detected using ABRicate/PlasmidFinder and minimum nucleotide identity and coverage thresholds of 80 and 60%, respectively; blue); (iii) presence and absence of antimicrobial resistance (AMR) and stress response genes (identified using AMRFinderPlus and default parameters; green). Core SNPs were identified using Snippy. The phylogeny was constructed using the results of ten independent runs using a relaxed lognormal clock model, the Standard_TPM1 nucleotide substitution model, and the Coalescent Bayesian Skyline population model implemented in BEAST version 2.5.1, with 10% burn-in applied to each run. LogCombiner-2 was used to combine BEAST 2 log files, and TreeAnnotator-2 was used to construct the phylogeny using common ancestor node heights. Abbreviations for the 15 antimicrobials are: AMK, amikacin; AMC, amoxicillin-clavulanic acid; AMP, ampicillin; FOX, cefoxitin; TIO, ceftiofur; CRO, ceftriaxone; CHL, chloramphenicol; CIP, ciprofloxacin; GEN, gentamicin; KAN, kanamycin; NAL, nalidixic acid; STR, streptomycin; SXT, sulfamethoxazole-trimethoprim; SUL, sulfisoxazole; TET, tetracycline.

Isolates from Farm 25 were partitioned into two clades: one containing *S.* Typhimurium isolates, and one containing *S.* Typhimurium Copenhagen isolates (based on SeqSero2’s *in silico* serotype assignments; Figure 10). Farm 25 isolates assigned to the *S.* Typhimurium Copenhagen variant (i) shared a common ancestor that existed circa December 2007/January 2008 (node age 2007.99, CA node height 95% HPD interval 2007.68-2008.21); (ii) were all resistant to ampicillin, kanamycin, streptomycin, sulfisoxazole, and tetracycline, with reduced susceptibility to additional antimicrobials observed sporadically; (iii) all possessed replicons for IncA/C2, IncFIB(AP001918), and IncFII(s) plasmids; and (iv) all possessed streptomycin resistance genes *aadA12*, *APH(3’’)-Ib* and *APH(6)-Id* (i.e., *strAB*), beta-lactamase *bla*_TEM-1_, and antiseptic resistance gene *qacE* delta 1, with other AMR/stress response genes present sporadically (Figure 10 and Supplementary Figure S10). Farm 25 isolates assigned to the *S.* Typhimurium clade shared a common ancestor that existed circa July 2007 (node age 2007.55, CA node height 95% HPD interval 2007.18-2007.78; Figure 10 and Supplementary Figure S10). All Farm 25 *S.* Typhimurium isolates were resistant to cefoxitin; resistance to additional antimicrobials, along with presence of IncI1 plasmids and *bla*_CMY-2_, was observed sporadically (Figure 10).

The isolate from Farm 17 was predicted to belong to the *S.* Typhimurium Copenhagen serotype variant using SeqSero2 and shared a common ancestor with the *S.* Typhimurium isolates from Farm 22, which existed circa 2000 (node age 1999.92, CA node height 95% HPD interval 1991.72-2006.21; Figure 10 and Supplementary Figure S10). Of the ten *S.* Typhimurium strains from Farm 22, seven were pan-susceptible (Figure 10). A bovine strain (BOV_TYPH_22_03−14−08_R8−0865) was resistant to ampicillin, ceftiofur, and ceftriaxone and was found to harbor IncI1 and IncI2 plasmids, as well as beta-lactamase *bla*_CTX-M-55_ (Figure 10). The remaining two bovine isolates were intermediately resistant to chloramphenicol and additionally resistant to either (i) amoxicillin-clavulanic acid, ampicillin, and sulfisoxazole, or (ii) tetracycline (Figure 10). Overall, isolates from Farm 17 and Farm 22 shared a common ancestor with the Farm 25 *S.* Typhimurium clade that existed circa 1988 (node age 1988.02, CA node height 95% HPD interval 1969.24-2002.47; Figure 10 and Supplementary Figure S10).

## 4 Discussion

### 4.1 WGS can be used to monitor pathogen microevolution and temporal AMR dynamics in animal reservoirs

Cattle may act as a reservoir for *Salmonella* and may facilitate its transmission to other animals (Mentaberre et al., 2013; Wiethoelter et al., 2015) or humans, either through direct contact or via the food supply chain (Hoelzer et al., 2011; Cummings et al., 2012; Mughini-Gras et al., 2014; An et al., 2017; Gutema et al., 2019). Even outside of a bovine host, *Salmonella* can survive in the farm environment for a prolonged amount of time, making persistent strains a particularly relevant threat to animal and human health (Rodriguez et al., 2006; Cummings et al., 2010b; Gorski et al., 2011; Toth et al., 2011; Tassinari et al., 2019). This threat can be compounded when persistent strains are exposed to antimicrobials, as a number of studies have linked antimicrobial exposure to the emergence of AMR in different foodborne pathogens, including *Salmonella*, *Escherichia coli*, and *Campylobacter* (Boerlin et al., 2001; McDermott et al., 2002; Delsol et al., 2003; Dutil et al., 2010; Hoelzer et al., 2017).

However, AMR acquisition among pathogens in livestock environments is far from absolute; in the absence of selective pressures (e.g., antimicrobial exposure), some AMR traits may be associated with a fitness cost for a given organism (Melnyk et al., 2015; Hoelzer et al., 2017; San Millan and MacLean, 2017). Consequently, interventions or changes in farm management practices (e.g., limiting antimicrobial use for all or selected antimicrobials, targeted use of some antimicrobials) may lead to reduced selection of AMR bacteria (Aarestrup, 2015; Tang et al., 2017; Scott et al., 2018). As such, the dynamics of AMR acquisition and loss among livestock-associated bacterial pathogens are complex and influenced by a wide range of factors, including the antimicrobials and treatment regimens used, farm management practices, environmental conditions, and the biology of the pathogens themselves (Aarestrup, 2015; Hoelzer et al., 2017; Davidson et al., 2018; Pereira et al., 2019; Clarke et al., 2020).

Using a WGS-based approach applied to serially sampled *Salmonella* strains isolated over a short time frame (i.e., less than two years), the study detailed here reveals that sporadic acquisition and loss of acquired AMR genes can occur within closely related populations over a short timescale. One particularly notable observation is represented by multiple, independent acquisitions of the beta-lactamase *bla*_CMY_ among *S.* Typhimurium and *S.* Typhimurium Copenhagen, as all *bla*_CMY_ acquisition events within this serotype group were confined to the 2000s. *bla*_CMY_ can confer resistance to cephalosporins, including (i) ceftriaxone, which has been used in human medicine since the early 1980s, and is used to treat invasive salmonellosis cases when fluoroquinolones cannot be used (e.g., for pediatric salmonellosis cases), and (ii) ceftiofur, which has been used in veterinary settings since the late 1980s to treat disease cases among dairy cattle and other animals (Hornish and Kotarski, 2002; Alcaine et al., 2005; Liebana et al., 2013; Yang et al., 2016; Carroll et al., 2017b; Carroll et al., 2020a). Because *bla*_CMY_ often confers resistance to both ceftriaxone and ceftiofur, there has been concern that the use of ceftiofur in livestock can contribute to the dissemination of *bla*_CMY_ and thus yield bacterial populations that are co-resistant to ceftriaxone (Alcaine et al., 2005; Tragesser et al., 2006; Carroll et al., 2017b; Carroll et al., 2020a).

Two independent *bla*_CTX-M_ acquisition events among *S.* Melegridis and *S.* Typhimurium were additionally observed. *bla*_CTX-M_, which also confers resistance to cephalosporins, was rarely detected in the United States in the 1990s (Lewis et al., 2007; Canton et al., 2012). However, *bla*_CTX-M_ rapidly increased in prevalence in the United States between 2000 and 2005 (Lewis et al., 2007; Canton et al., 2012), and there is evidence that bacterial populations associated with dairy cattle may have been affected as well. In a study of *E. coli* isolated from dairy cattle in the western United States, the prevalence of *bla*_CTX-M_ was found to have increased between 2008 and 2012 (Afema et al., 2018). The results of our study are congruent with these findings, as all observed *bla*_CTX-M_ acquisition events were estimated to have occurred in the 2000s.

AMR loss events were additionally observed among the bovine-associated, MDR *S.* Newport isolates sequenced here. Prevalence of MDR *S.* Newport among humans increased rapidly in the United States within the late 1990s and early 2000s and was linked to cattle exposure, farm/petting zoo exposure, unpasteurized milk consumption, and ground beef consumption (Spika et al., 1987; Gupta et al., 2003; Karon et al., 2007). While chloramphenicol resistance is often a hallmark characteristic of MDR *S.* Newport, the MDR *S.* Newport lineage represented by an isolate in this study was chloramphenicol-susceptible and was predicted to have lost chloramphenicol resistance gene *floR* after 2008. These results indicate that even well-established MDR pathogens can still be subjected to temporal changes in AMR profile.

Due to the global burden that AMR pathogens impose on the health of humans and animals, numerous agencies have called for improved monitoring of pathogens and their associated AMR determinants along the food supply chain (World Health Organization, 2014; 2017; Centers for Disease Control and Prevention, 2019). The study detailed here showcases how WGS can be used to identify temporal changes in the resistomes of livestock-associated pathogens at the farm level. However, further sequencing efforts querying (i) a larger selection of *Salmonella* strains isolated from livestock on individual farms (ii) over a longer timeframe are needed to determine whether the AMR dynamics observed here are merely sporadic, or rather are indications of larger trends.

### 4.2 Bovine-associated *Salmonella* lineages with heterogeneous antimicrobial resistance profiles may be present across multiple farms or strongly farm-associated

Geography has been shown to play an important role in shaping bacterial populations (Achtman, 2008; Strachan et al., 2015), including some *Salmonella* lineages (Carroll et al., 2017b; Palma et al., 2018; Fenske et al., 2019; Liao et al., 2020). However, for some foodborne pathogens, including some *Salmonella* populations, global spread of lineages due to human migration and movement of food and animals can often obfuscate local phylogeographic signals (Wong et al., 2015; Llarena et al., 2016; The et al., 2016; Palma et al., 2018).

In the study detailed here, *Salmonella* lineages isolated from cattle and their associated environments on 13 separate farms in a confined geographic location (i.e., New York State) were found to vary in terms of the farm-specific signal they possessed; some lineages (i.e., *S.* Anatum, *S.* Newport, *S.* Typhimurium, *S.* Typhimurium Copenhagen, some *S.* Kentucky populations) were found to be strongly associated with a particular farm, while other lineages (i.e., *S.* Cerro, *S.* Meleagridis, some *S.* Kentucky populations) were distributed across multiple farms. Multiple scenarios may explain the existence of *Salmonella* lineages distributed across multiple farms, including movement of livestock, humans, pets, and/or wildlife (Skov et al., 2008; Hoelzer et al., 2011; Palma et al., 2018) or introduction via feed; however, additional metadata (e.g., farm geography, proximity to other farms in the study, and management practices) are needed to draw further conclusions. Even with limited metadata available, WGS data can provide important insights into *Salmonella* transmission and introduction on farms, as shown in this study. For example, for one farm (i.e., Farm 25), two *Salmonella* Typhimurium clonal groups were present (i.e., one representing Typhimurium and one representing Typhimurium Copenhagen), each of which shared a common ancestor dated circa 2007. WGS data can be used to identify time frames in which *Salmonella* lineages may have emerged in a given farm or region, which could help pinpoint root causes (e.g., changes in management practices that occurred around the predicted time of emergence).

While the characterization of additional, larger strain sets from more geographically diverse farms is essential, our data suggest that specific *Salmonella* clones may persist on a given farm. This suggests that WGS databases covering isolates from a large number of farms could be used to develop initial hypotheses about farm sources of *Salmonella* strains. While such applications are tempting, it is crucial that these types of data are only used for initial hypothesis generation; rigorous, critical epidemiological investigations are essential before any conclusions regarding strain source are drawn.

### 4.3 *In silico* serotyping of bovine-associated *Salmonella* can outperform traditional serotyping

Well into the genomic era, serotyping remains a vital microbiological assay that allows *Salmonella* isolates to be classified into meaningful, evolutionary units. Serotype assignments are used to facilitate outbreak investigations and surveillance efforts, construct salmonellosis risk assessment frameworks, and inform food safety and public health policy and decision-making efforts (Yoshida et al., 2016; Gutema et al., 2019). Importantly, serotyping is used worldwide to monitor salmonellosis cases among humans and animals, including cattle (Gutema et al., 2019; Centers for Disease Control and Prevention, 2020).

In this study, serotypes assigned using traditional phenotypic methods were compared to serotypes assigned using two *in silico* methods (i.e., SISTR and SeqSero2). Notably, both *in silico* serotyping approaches outperformed traditional *Salmonella* serotyping for this data set. Serotypes assigned using SISTR’s cgMLST approach and/or SeqSero2 were congruent with the *Salmonella* whole-genome phylogeny and were able to resolve all un-typable, ambiguous, and incorrectly assigned serotypes (Supplementary Table S1). It is essential to note that the data set queried here is far too small and, thus, inadequate to formally benchmark these tools. Furthermore, all serotypes studied here were among the ten most frequently reported serotypes of *Salmonella* isolated from subclinical cattle between 2000 and 2017 (Gutema et al., 2019), indicating that they are well-represented in public databases and thus likely do not pose a significant challenge to *in silico* tools. However, the results observed here reflect results observed in several recent studies querying larger numbers of isolate genomes and/or a wider array of diverse *Salmonella* serotypes (Yachison et al., 2017; Ibrahim and Morin, 2018; Diep et al., 2019; Banerji et al., 2020; Uelze et al., 2020). In their analysis of 1,624 animal- and food-associated (i.e., non-human) *Salmonella* isolate genomes assigned to 72 serotypes, Uelze et al. (Uelze et al., 2020) reported that SISTR and SeqSero2 achieved the highest and second-highest accuracy of all tested *in silico Salmonella* serotype assignment tools, correctly serotyping 94% and 87% of isolates, respectively. However, unlike the results observed here, the authors note that neither tool outperformed traditional serotyping conducted by *Salmonella* reference laboratories. Similarly, in a study of 813 *Salmonella* isolates, SISTR outperformed the original version of SeqSero (i.e., SeqSero 1.0) with serotype prediction accuracies of 94.8% and 88.2%, respectively (Yachison et al., 2017).

With WGS data in hand, *in silico* serotyping is rapid, scalable, inexpensive, and reproducible (Uelze et al., 2020). Nevertheless, it is important to be mindful of the strengths and weaknesses of different *in silico* serotyping tools. In their benchmarking study, Uelze et al. (Uelze et al., 2020) recommended SISTR as the optimal contemporary tool for routine *in silico Salmonella* serotyping based on overall accuracy; however, they additionally report that the raw read mapping approach implemented in SeqSero2 (i.e., “allele mode”) outperforms SISTR for prediction of monophasic variants. Banerji et al. (Banerji et al., 2020) did not assess the performance of SISTR on their data set, as it requires assembled genomes and not raw reads (another potential drawback if a high-quality assembly is not available or obtainable for an isolate of interest); however, they found that both SeqSero and MLST approaches misidentified monophasic variants, particularly among the important monophasic *S.* Typhimurium lineage. Among the bovine-associated *Salmonella* strains sequenced here, we observed a combination of *S.* Typhimurium strains that possessed the O5 epitope, and those that did not (i.e., *S.* Typhimurium Copenhagen). Importantly, SISTR was unable to differentiate *S.* Typhimurium from *S.* Typhimurium Copenhagen, while SeqSero2 could, as reported previously (Ibrahim and Morin, 2018; Zhang et al., 2019). While the differentiation of *S.* Typhimurium from its O5-counterpart may not be essential for all microbiological applications, it is important to be aware of this limitation; *S.* Typhimurium Copenhagen has been responsible for outbreaks and illnesses around the world (Luceron et al., 2017; Tack et al., 2020) and can be multidrug-resistant, as demonstrated here and elsewhere (Frech et al., 2003; Tack et al., 2020).

While serotypes assigned *in silico* using SISTR and SeqSero2 are highly accurate and congruent, each tool has strengths and limitations; as such, an approach that utilizes both methods, such as the one employed here, may increase accuracy and minimize potential misclassifications. Results from other studies support this (Yachison et al., 2017; Banerji et al., 2020). For example, in an analysis of 520 primarily human-associated *Salmonella* isolate genomes, Banerji et al. (Banerji et al., 2020) found that serotypes assigned *in silico* using SeqSero showed 98% concordance with traditional serotyping and outperformed serotype assignment using seven-gene MLST. However, when SeqSero and seven-gene MLST were used in combination, *in silico* serotyping accuracy surpassed 99%, consistent with our results that a combination of SeqSero2 and cgMLST-based serotyping (as implemented in SISTR) improved *in silico* serotyping accuracy. Overall, the results provided here lend further support to the idea that *in silico* serotyping may eventually replace traditional serotyping as WGS becomes more widely used and accessible (Yachison et al., 2017; Banerji et al., 2020; Uelze et al., 2020).

### 4.4 Limitations of the *in silico* AMR method evaluation presented here, and considerations for future AMR monitoring efforts among livestock and beyond

In addition to studying the microevolution and AMR dynamics of bovine *Salmonella* on a genomic scale, the study presented here compared results obtained from numerous *in silico* AMR characterization pipelines that attempt to replicate traditional microbiological assays used to characterize AMR *Salmonella*. More specifically, each of the following tools was applied to the set of 128 bovine-associated *Salmonella* genomes sequenced here: (i) combinations of five *in silico* AMR determinant detection pipelines (i.e., ABRicate, AMRFinderPlus, ARIBA, BTyper, and SRST2) and one to five AMR determinant databases (i.e., ARG-ANNOT, CARD, MEGARes, NCBI, and ResFinder); and (ii) an *in silico* MIC prediction tool (i.e., PATRIC3).

Here, all AMR determinant detection pipelines and AMR determinant databases showed an extremely high degree of concordance; regardless of pipeline or database selection, all tools performed nearly identically on an SIR-prediction task relative to (i) “true” SIR classifications based on NARMS breakpoints and “true” MIC values obtained for a panel of 15 antimicrobials, and (ii) each other. A previous, small-scale (*n* = 111) WGS-based study of AMR *Salmonella* observed similarly high rates of concordance among several *in silico* AMR determinant detection tools (Cooper et al., 2020). However, in addition to its small sample size, this study also relied on *Salmonella* strains isolated from a single source (broiler chickens) in a single country (Canada) over an extremely short temporal range (December 2012-December 2013). Similarly, the study detailed here is not a formal benchmarking study, and it is essential that its numerous limitations are pointed out.

First and foremost, the study conducted here relied on WGS data from an extremely small sample of *Salmonella* isolates (*n* = 128) from a single source (dairy cattle and their surrounding farm environments) in a confined geographic area (New York State, United States) isolated over a short temporal range (fewer than two years). While all isolates were “unique” (i.e., each strain was isolated from a separate sampling event of a unique source), many isolates were highly similar at both the genomic and pan-genomic level (e.g., *S.* Cerro, *S.* Meleagridis), indicating that, in some cases, the same lineage was being sampled repeatedly over time. Consequently, this relatively miniscule sample is unrepresentative of AMR pathogens and, more specifically, *Salmonella* as a whole; readers should not infer the general superiority or inferiority of any AMR detection tool or database tested, and the results obtained here should not be extrapolated to external data sets.

Secondly, the data set queried here was heavily biased towards susceptible isolates. More than half of all isolates were pan-susceptible to the 15 antimicrobials included on the panel, and only 21 unique phenotypic SIR profiles were observed. Congruent with this, relatively little diversity was observed in terms of AMR gene profile (e.g., the AMRFinderPlus pipeline produced 20 unique AMR/stress response determinant presence/absence profiles among the 128 isolates sequenced here). This is not particularly surprising; numerous studies have shown that the resistomes of bovine-associated *Salmonella* tend to be less diverse than *Salmonella* isolated from humans (Afema et al., 2015; Carroll et al., 2017b), as well as some other animals (Mellor et al., 2019). Furthermore, the resistomes of *Salmonella* isolated from subclinical cattle, such as the isolates queried in this study, have been shown to be less diverse than the resistomes of *Salmonella* isolated from cattle showing clinical signs of disease (Afema et al., 2015).

The relative homogeneity of the subclinical *Salmonella* bovine resistome and bias towards antimicrobial-susceptible isolates have important implications for the AMR pipeline/database comparison conducted here. For this data set, stringent and conservative approaches are rewarded, as isolates that do not possess AMR determinants are more likely to be predicted to be susceptible. While it is possible that different AMR detection tools may perform better on WGS data from pathogens with more diverse resistomes, very few formal benchmarking studies of *in silico* AMR determinant detection tools currently exist (Hendriksen et al., 2019). The choice of AMR determinant database in combination with the choice of pipeline, on the other hand, can clearly affect AMR determinant identification in a critical way. For example, the ARG-ANNOT database (Gupta et al., 2014), a manually curated AMR determinant database first published in 2014, is not updated as frequently as other AMR databases (e.g., CARD, NCBI, ResFinder). Since it was last updated in May 2018 (accessed May 25, 2020), at the time of our study, the database does not yet include three novel plasmid-mediated genes (*mcr-8*, *-9,* and *-10*) that can confer resistance to colistin, a last-resort antibiotic used to treat MDR and extensively drug resistant infections (Wang et al., 2018; Carroll et al., 2019; Wang et al., 2020). Similarly, versions of tools that rely on even older versions of this database would not be able to detect all members of the continuously growing repertoire of *mcr* genes. For the low-diversity subclinical bovine *Salmonella* resistomes queried here, the use of a smaller database was inconsequential, as reflected in the high congruency of all methods and databases observed here. For some studies, a manually curated database of AMR genes that is updated conservatively may possibly even be desirable, as such a database may yield less noise and improve interpretability and reproducibility. However, for pathogens with more diverse resistomes (e.g., human clinical isolates, isolates from geographic regions with different antibiotic use practices), the omission of critically important genes could be a disastrous flaw. Similar to our results, a large study (*n* = 6,242) querying NARMS isolates belonging primarily to the *Salmonella enterica* species (*n* = 5,425) observed a high degree of concordance between NCBI’s AMRFinder tool and ResFinder (this study was used to validate AMRFinder and the NCBI AMR determinant database) (Feldgarden et al., 2019). However, when differences between tools were observed, the vast majority (81%) were attributed to differences in database composition (Feldgarden et al., 2019; Hendriksen et al., 2019).

Thirdly, the small sample size (*n* = 128) and sparsity of AMR isolates available in this study limited the methods that could be used to formulate the AMR tool/database comparison. The approach used here was similar to the one used to validate the AMRFinder tool (Feldgarden et al., 2019) in that it relied on known AMR-determinant/AMR phenotype associations available in the literature (see Supplemental Table S4 of Feldgarden, et al.) (Feldgarden et al., 2019). As such, the approach used here does not account for previously unobserved genotype/phenotype associations. Furthermore, different variants of the same AMR gene may yield different AMR phenotypes; for example, some variants of the OXA beta-lactamases are able to confer resistance to cephalosporins, while others are not (Evans and Amyes, 2014). All AMR determinant detection pipelines tested here produced nearly identical genes calls among the 128 isolates sequenced here, and all detected AMR determinants were manually annotated in a consistent fashion; while the overall accuracy of all pipeline/database combinations tested here could likely be optimized if more accurate, data set-specific AMR genotype/phenotype associations could be derived, congruency between tools/databases would likely remain high.

Classifying bacterial pathogens into discrete SIR groups using AMR determinant detection methods is challenging, as it requires users to have a great degree of prior knowledge regarding the AMR determinants that are detected, the antimicrobials of interest, and the pathogen being studied. PATRIC3’s MIC prediction tool (Nguyen et al., 2019) offers a promising departure from this framework, as it allows for the prediction of MIC values directly from WGS data. Interpreting the resulting *in silico* MIC values does not require any prior knowledge on the user end, and results can be harmoniously integrated into the SIR framework using clinical breakpoints. Among the *Salmonella* isolates sequenced here, SIR classification using PATRIC3 resulted in an overall accuracy of 93%. However, all of the limitations of this study described above for AMR determinant detection (e.g., small sample size, AMR sparsity bias, single-source, single geographic region, small temporal range) apply to *in silico* MIC prediction as well; for example, when “intermediate” resistance predictions produced via PATRIC3 are re-classified as “susceptible” (as was done for the AMR determinant detection approaches used here), PATRIC3’s accuracy for this data set increases to 96.0% and is on par with all other AMR prediction methods tested here. Readers should thus interpret comparisons between these methods with caution.

Benchmarking and validating AMR detection and prediction tools is notoriously challenging (Feldgarden et al., 2019), and very few researchers have undertaken this task (Hendriksen et al., 2019). While high congruency may be observed between tools (Clausen et al., 2016), identification of a clear “optimal” method for *in silico* AMR characterization has remained elusive, as the few available benchmarking studies differ in terms of the tools tested, the AMR database(s) used, and the data set(s) chosen for benchmarking. Furthermore, the underlying WGS data can affect pipeline performance (Clausen et al., 2016; Feldgarden et al., 2019). For example, assembly quality has been shown to influence AMR determinant detection for methods that rely on assembled genomes (Clausen et al., 2016; Hendriksen et al., 2018; Hendriksen et al., 2019). Thus, whether a read- or assembly-based method performs optimally can depend on a given data set (e.g., sequencing quality, the organism being studied). Another criticism of BLAST-based AMR gene detection methods among assembled genomes has been the choice of thresholds used for considering AMR determinants present or absent (Hendriksen et al., 2019). Here, no significant differences were observed between the accuracy of read-based ARIBA and SRST2 and assembly-based ABRicate, AMRFinderPlus, and BTyper. Additionally, for BLAST-based methods, a relatively wide range of optimal nucleotide identity and coverage values were found to maximize accuracy, with thresholds of 75% identity and 50-60% coverage adequate for most pipeline/database combinations. Overall, when selecting an *in silico* AMR characterization method, researchers should take into account not only practical considerations (e.g., whether reads or assembled genomes are available, the quality of reads and/or assembled genomes), but also the biology of the pathogen being studied (e.g., by querying organism-specific, AMR-conferring point mutations). To assess the robustness of *in silico* AMR predictions, researchers may additionally consider employing multiple *in silico* AMR characterization tools and/or databases in combination, as well as testing various AMR gene detection thresholds.

Finally, it is essential to note that accuracy estimates for *in silico* AMR characterization tools relative to gold-standard phenotypic methods are only as reliable as the phenotypic data they rely on. Previous studies of *Salmonella* that attempted to predict phenotypic AMR using *in silico* methods (McDermott et al., 2016; Cooper et al., 2020) have reported accuracy values between 98 and 100%. For this data set, the highest accuracy achieved was 97.4% (for SRST2/ARG-ANNOT). However, sensitivity (i.e., the ability of an *in silico* pipeline/database combination to correctly classify an isolate as phenotypically resistant to an antimicrobial) was lower for this study (71.8-84.4%) than sensitivity estimates calculated in a study of MDR *Salmonella* that included bovine isolates from New York State (97.2%) (Carroll et al., 2017b). As mentioned above, this could be due to the sparsity of AMR among isolates in this data set (i.e., predicting susceptible, rather than resistant phenotypes, is incentivized here). However, it is important to note that the AMR phenotypes of several isolates were highly incongruent with their respective AMR genotypes, regardless of the tool/database used for *in silico* AMR prediction. For example, one *S.* Cerro isolate (BOV_CERO_35_10-02-08_R8-2685) was reported to be phenotypically resistant to nine antimicrobials but did not possess any acquired AMR determinants known to produce this phenotypic AMR profile. A recent case study in which WGS and phenotypic methods were used to characterize *Salmonella* isolates from raw chicken identified numerous AMR genotype/phenotype discrepancies resulting in both false negative and false positive predictions for *in silico* methods (Zwe et al., 2020). In this case study, the authors attributed these discrepancies to heteroresistant *Salmonella* subpopulations (i.e., a subpopulation of bacteria that exhibits a range of susceptibility to a particular antimicrobial). The possibility that several heteroresistant *Salmonella* populations were characterized here cannot be discounted, as isolates underwent phenotypic AMR characterization and WGS separately (i.e., years apart). Other biological phenomena, such as plasmid loss during storage or culturing, or unknown/undetected resistance genes or mutations, could also contribute to discrepancies (Hendriksen et al., 2018). However, it is also possible that one or more incongruent isolates was mislabeled and/or mishandled during AMR phenotyping, genomic DNA extraction, and/or WGS. While removal of these isolates from the data set would increase overall prediction accuracy, the high congruency between AMR genotyping methods would be unaffected.

## 5 Author Contributions

LMC designed and carried all computational analyses. AJB, AG, JDS, KJC, and RAC collected, analyzed, and/or interpreted all microbiological data. LMC and MW conceived the study and co-wrote the manuscript with input from all authors.

## 6 Funding

This material is based on work supported by the National Science Foundation Graduate Research Fellowship Program under grant no. DGE-1650441.

## 7 Conflict of Interest

The authors declare that the research was conducted in the absence of any commercial or financial relationships that could be construed as a potential conflict of interest.

## 8 Acknowledgments

This project was supported in part by the Cornell University Zoonoses Research Units of the Food and Waterborne Diseases Integrated Research Network, funded by the National Institute of Allergy and Infectious Diseases, National Institutes of Health, under contract no. N01-AI-30054.

## Notes

### Competing Interest Statement

The authors have declared no competing interest.

