## Supplementary Figure S1 for "Monitoring the Antimicrobial Resistance Dynamics of *Salmonella enterica* in Healthy Dairy Cattle Populations at the Individual Farm Level Using Whole-Genome Sequencing"

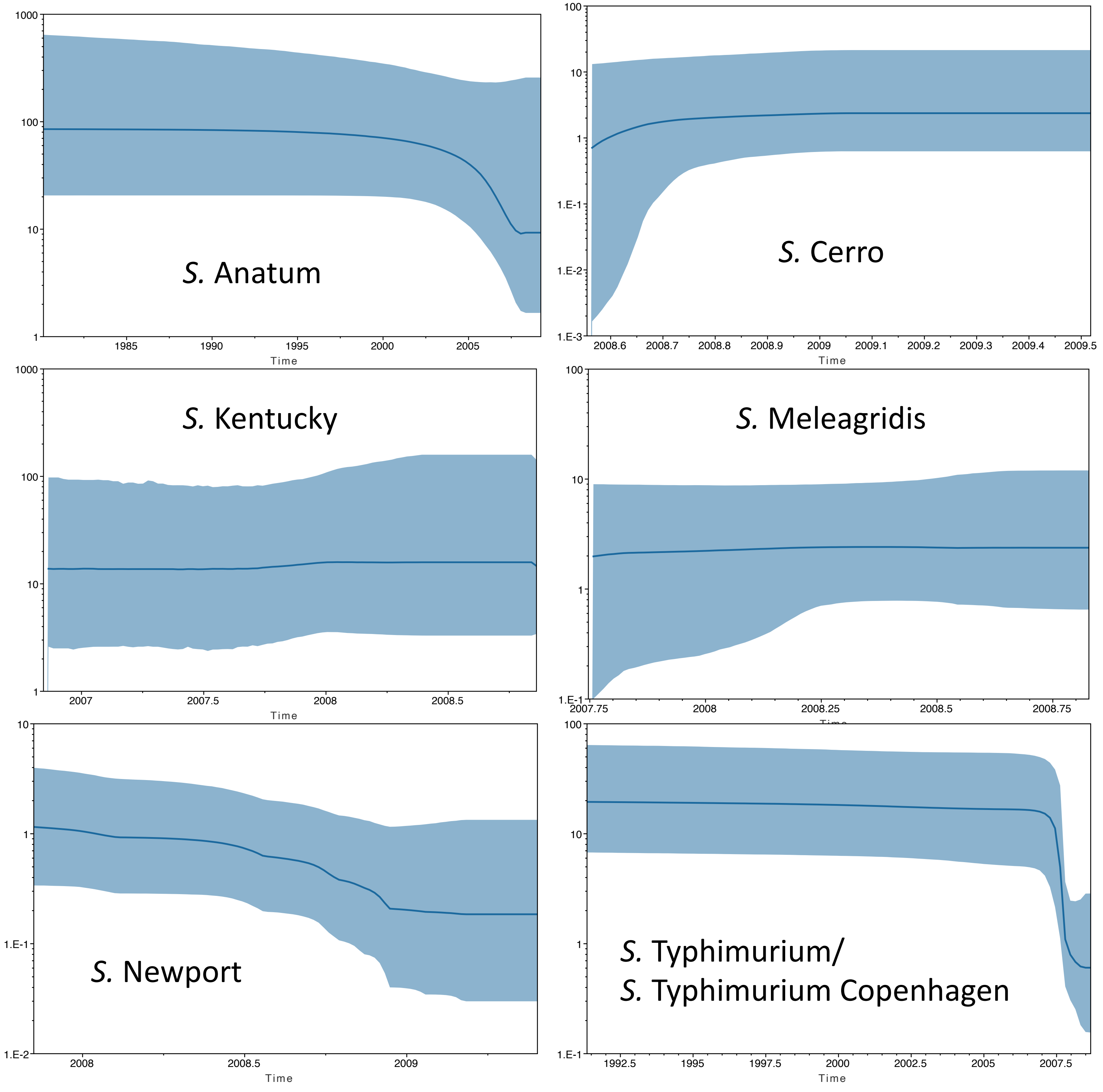


**Supplementary Figure S1.** Skyline plots constructed for each *Salmonella* serotype group. Effective population size and time in years are plotted on the Y- and X-axes, respectively. The median effective population size estimate is denoted by the blue line, with upper and lower 95% highest posterior density interval bounds denoted by blue shading.
