## Supplementary Figure S2 for "Monitoring the Antimicrobial Resistance Dynamics of *Salmonella enterica* in Healthy Dairy Cattle Populations at the Individual Farm Level Using Whole-Genome Sequencing"

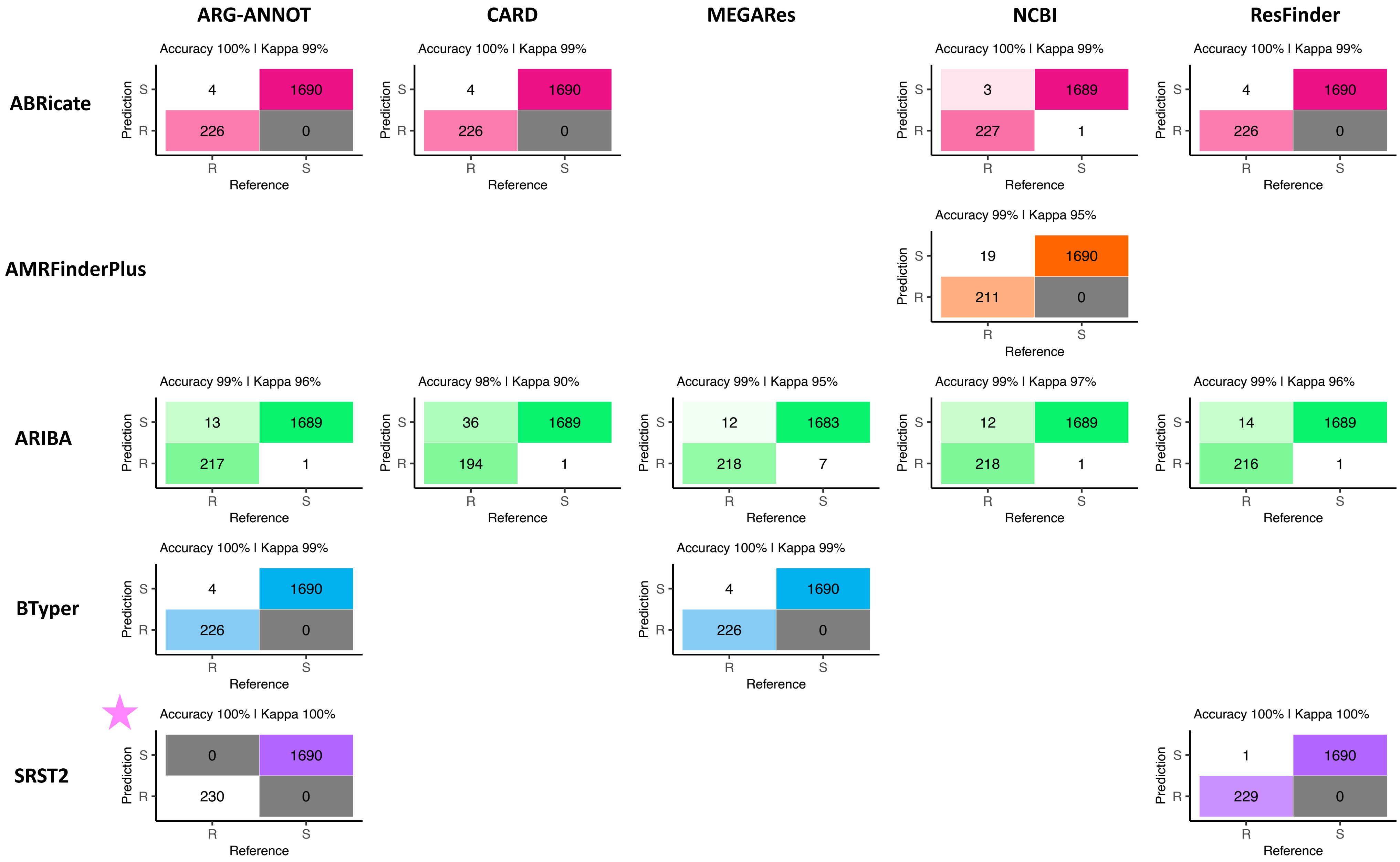


**Supplementary Figure S2.** Confusion matrices showcasing agreement between susceptible/resistant (S and R, respectively) classification of 128 *Salmonella* isolates obtained using SRST2/ARG-ANNOT (denoted as the matrix "Reference") and other *in silico* methods (denoted as the matrix "Prediction") for 15 antimicrobials. "Accuracy" values above each matrix denote the percentage of instances classified identically to SRST2/ARG-ANNOT (annotated with a pink star), out of all instances (i.e., not necessarily the "true" phenotypic S/R classification). Kappa values above each matrix denote Cohen's kappa coefficient for the matrix, reported as a percent. Combinations of five pipelines (rows) and one to five AMR determinant databases (columns) were tested; isolate genomes that harbored one or more AMR determinants previously known to confer resistance to a particular antimicrobial were categorized as resistant to that antimicrobial ("R"), while those which did not were categorized as susceptible ("S"; see Supplementary Table S2 for all detected AMR determinants and their associated resistance classifications). For pipelines that relied on nucleotide BLAST (i.e., ABRicate and BTyper), the confusion matrix obtained using the nucleotide identity and coverage threshold combination that produced the highest accuracy relative to phenotypic data are shown (Supplementary Figure S3); for all other methods, confusion matrices obtained using default detection parameters are shown.
