## Supplementary Figure S3 for "Monitoring the Antimicrobial Resistance Dynamics of *Salmonella enterica* in Healthy Dairy Cattle Populations at the Individual Farm Level Using Whole-Genome Sequencing"

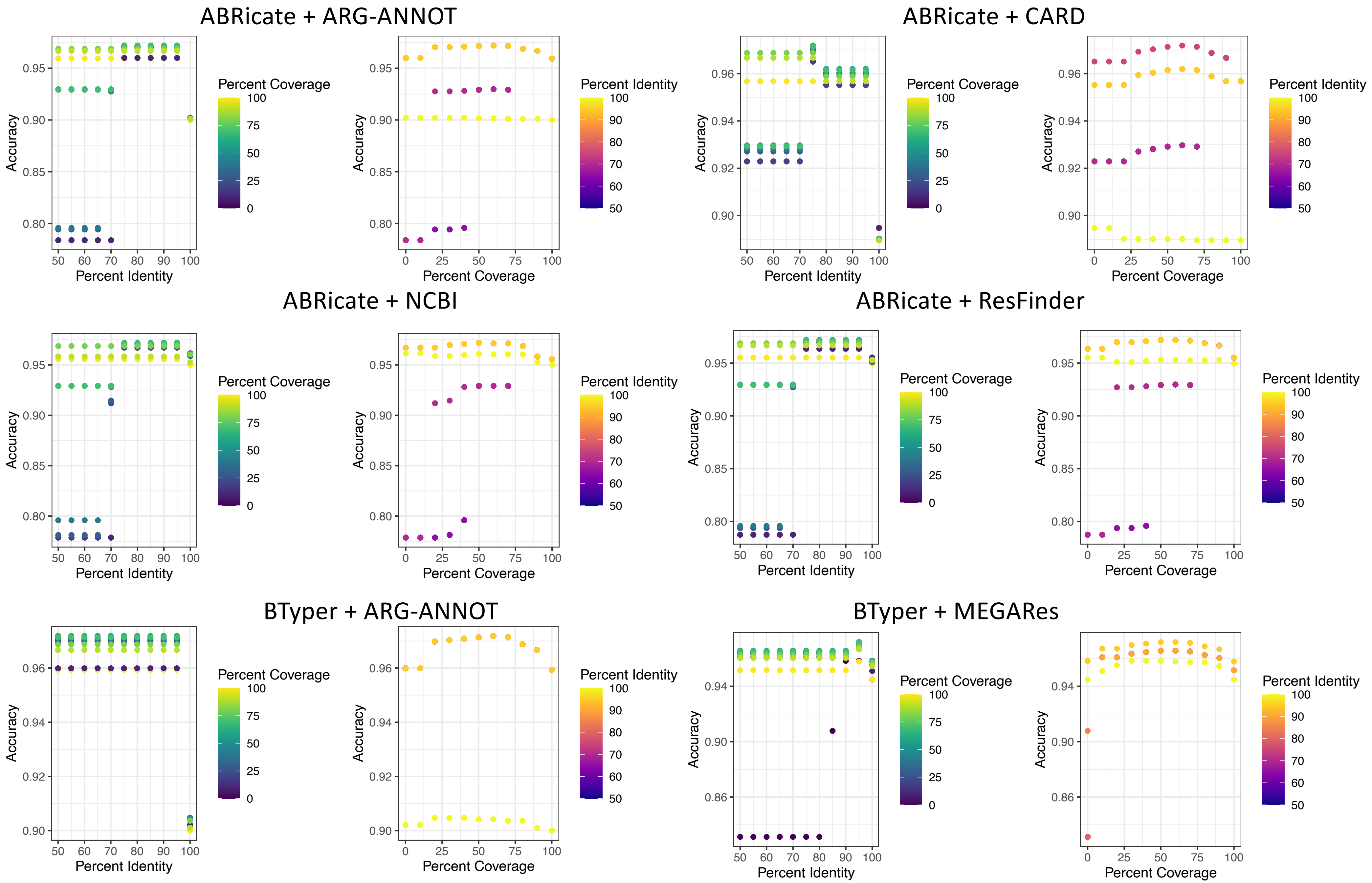


**Supplementary Figure S3.** Accuracy of nucleotide BLAST-based antimicrobial resistance (AMR) determinant detection methods (i.e., ABRicate and BTyper), using various database combinations (i.e., one of ARG-ANNOT, CARD, NCBI, MEGARes, or Resfinder), minimum percent nucleotide identity thresholds (i.e. "Percent Identity"), and minimum percent query coverage thresholds (i.e., "Percent Coverage"). Isolate genomes that harbored one or more AMR determinants previously known to confer resistance to a particular antimicrobial were categorized as resistant to that antimicrobial ("R"), while those which did not were categorized as susceptible ("S"; see Supplementary Table S2 for all detected AMR determinants and their associated resistance classifications). For all AMR determinant detection methods, isolates that showed intermediate phenotypic resistance to an antimicrobial were categorized as susceptible ("S") rather than resistant, as this classification produced slightly better accuracy scores for all pipeline/database combinations.
