## Supplementary Figure S4 for "Monitoring the Antimicrobial Resistance Dynamics of *Salmonella enterica* in Healthy Dairy Cattle Populations at the Individual Farm Level Using Whole-Genome Sequencing"

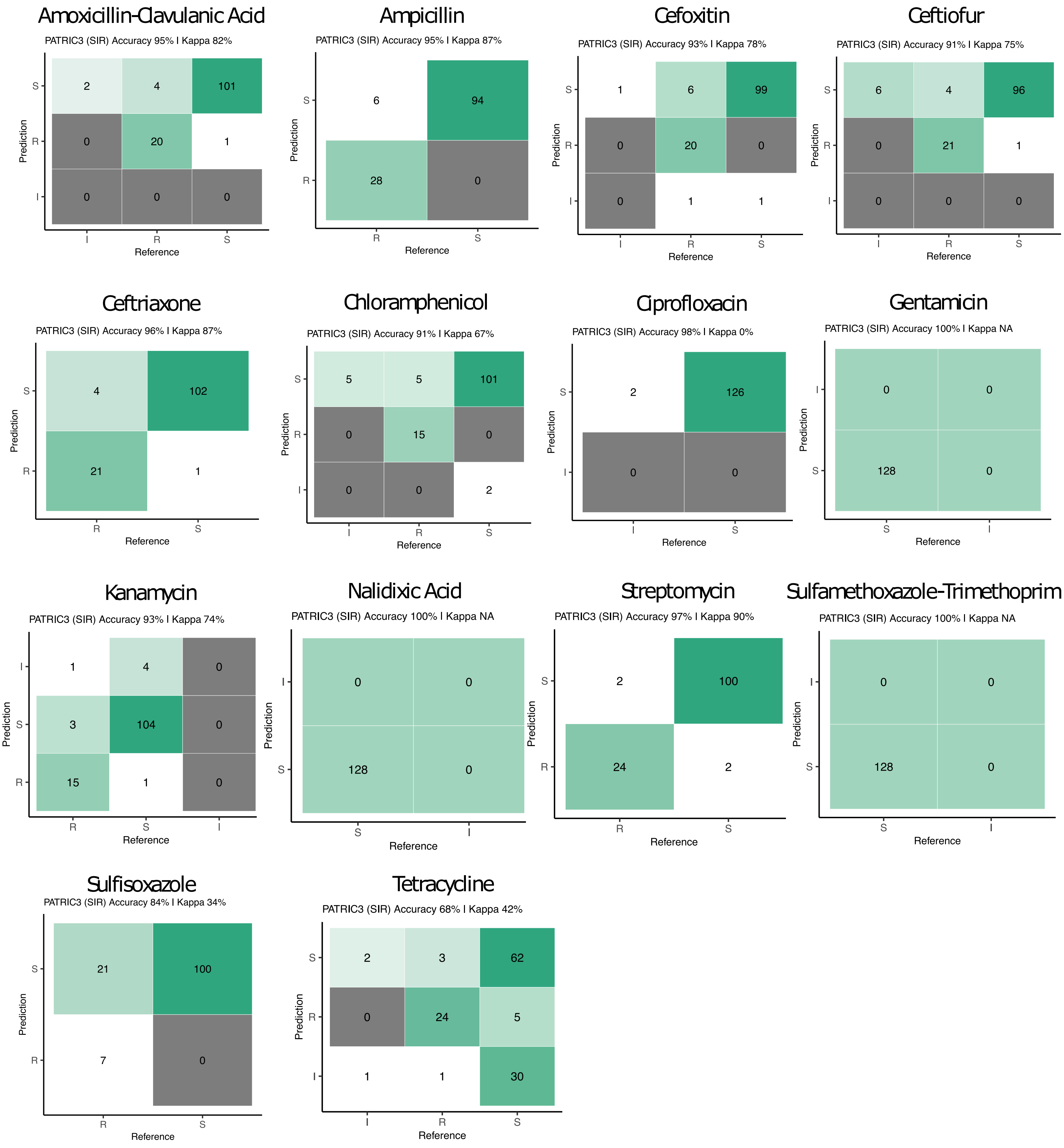


**Supplementary Figure S4.** Confusion matrices showcasing agreement between susceptible-intermediate-resistant (SIR) classification of all 128 *Salmonella* isolates obtained using phenotypic resistance testing (denoted as the matrix "Reference") and PATRIC3 (denoted as the matrix "Prediction") for 14 antimicrobials. The accuracy value above the matrix denotes the percentage of correctly classified instances out of all instances. The Kappa value above the matrix denotes Cohen's kappa coefficient for the matrix, reported as a percent. For both the phenotypic and PATRIC3 methods, SIR classification was determined using NARMS breakpoints for *Salmonella* (accessed March 23, 2020).
