## Supplementary Figure S10 for "Monitoring the Antimicrobial Resistance Dynamics of *Salmonella enterica* in Healthy Dairy Cattle Populations at the Individual Farm Level Using Whole-Genome Sequencing"

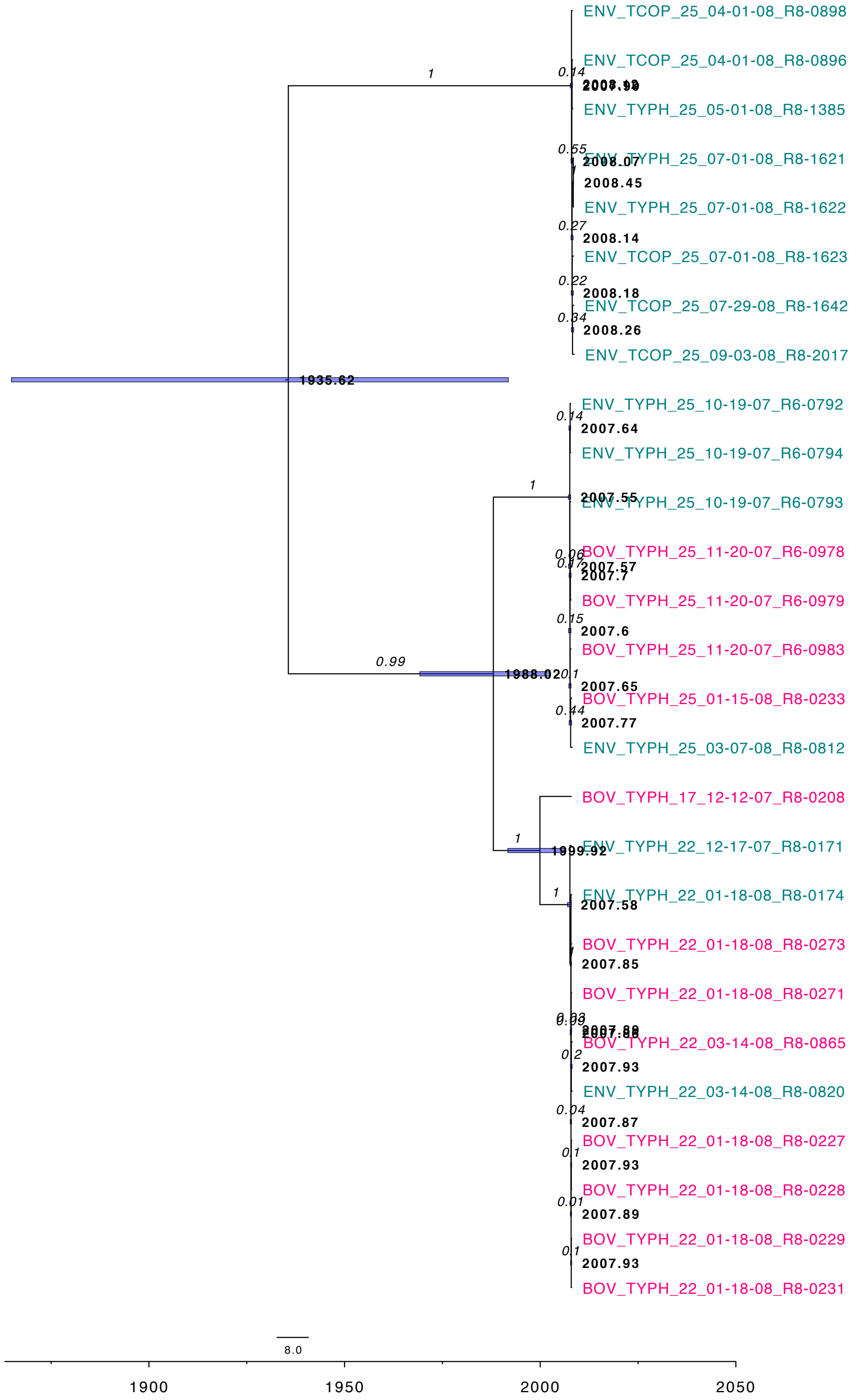


**Supplementary Figure S10.** Rooted, time-scaled maximum clade credibility (MCC) phylogeny constructed using core SNPs identified among 27 *Salmonella* Typhimurium and Typhimurium Copenhagen genomes isolated from subclinical bovine sources (pink tip labels) and the surrounding bovine farm environment (teal tip labels). Branch labels (black italic text) denote posterior probabilities of branch support, while node labels (black boldfaced text) denote node ages. Time in years is plotted along the X-axis, and branch lengths are reported in years. Node bars denote 95% highest posterior density (HPD) intervals for common ancestor (CA) node heights. Core SNPs were identified using Snippy. The phylogeny was constructed using the results of ten independent runs using a relaxed lognormal clock model, the Standard_TPM1 nucleotide substitution model, and the Coalescent Bayesian Skyline population model implemented in BEAST version 2.5.1, with 10% burn-in applied to each run. LogCombiner-2 was used to combine BEAST 2 log files, and TreeAnnotator-2 was used to construct the phylogeny using CA node heights.
